# Molecular basis of quorum-sensing signal transduction by CqsS and its inhibition by CqsA the autoinducer synthase

**DOI:** 10.64898/2026.05.05.722802

**Authors:** Mengchan Liu, Yufan Yang, Julie S. Valastyan, Yuru Zhang, Grace A. Beggs, Yanhong Liu, Zhenhua Wang, Bonnie L. Bassler, Hongwu Qian

**Affiliations:** Department of Cardiology, The First Affiliated Hospital of USTC, MOE Key Laboratory for Membraneless Organelles and Cellular Dynamics, Hefei National Research Center for Interdisciplinary Sciences at the Microscale, Division of Life Sciences and Medicine, University of Science and Technology of China, Hefei 230027, China; Department of Molecular Biology, Princeton University, Princeton, New Jersey, United States of America; Howard Hughes Medical Institute, Chevy Chase, Maryland, United States of America

## Abstract

Vibrio bacteria possess multiple quorum-sensing systems, each of which conveys information into the cell concerning the cell density and species composition of the vicinal community. Germane to the present work is the CqsA-CqsS quorum-sensing pathway that vibrios use for intra-genus communication. CqsA is the synthase for the autoinducer (*S*)-3-hydroxytridecan-4-one (CAI-1), and CqsS, a bi-functional two-component kinase-phosphatase, is the CAI-1 receptor. Here, we determine the cryo-EM structures of the full-length *Vibrio harveyi* CqsS dimer in the apo- and ATPγS-bound states. Our structures reveal the molecular basis for the initiation of the phosphorylation cascade, which we support through biochemical and genetic analyses. Autophosphorylation of CqsS occurs in *cis* at the conserved histidine residue (H1), whereas phospho-transfer to the conserved aspartate residue (D1) in the C-terminal receiver domain occurs in *trans*. By co-expressing CqsS and CqsA to capture CqsS bound to CAI-1, we unexpectedly discover a tight interaction between the CqsS intracellular domain and CqsA. In the complex, the CqsS H1 residues are positioned away from the catalytic cavity, and CqsS kinase activity is inhibited. Cryo-EM analysis, focused on the transmembrane domain of CqsS in this complex, reveals the presence of two CAI-1 molecules. CAI-1 binding induces an obvious movement of transmembrane helix 6, disrupting its extension into the cytosol, providing a structural basis for how CAI-1 binding in the periplasm is coupled to signal transduction into the cytoplasm. Together, our findings advance the understanding of CqsS-CAI-1-mediated quorum-sensing signal transduction, an activity that underpins the coordination of group behaviors in vibrios.

## Introduction

Quorum sensing (QS) is a cell-to-cell chemical communication process used by bacteria to orchestrate group behaviors^1^, including bioluminescence^2^, virulence factor production^3^, biofilm formation^4^, and DNA exchange^5^. QS relies on the production, release, and detection of extracellular signal molecules called autoinducers^6–8^.

Vibrios, the established model bacterial system for discoveries and analyses of QS-mediated communication, possess multiple QS systems. The systems are composed of autoinducer-receptor pairs that function in parallel to regulate group behaviors^8^. Important to the present work is the CqsS receptor that detects the cognate autoinducer CAI-1 ((*S*)-3-hydroxytridecan-4-one) and the CAI-1 synthase CqsA^9,10^. CqsS-CAI-1 fosters intra-genus cell-cell communication among vibrios^11^. Indeed, every sequenced vibrio species possesses CqsA and CqsS. CqsS is a membrane-bound two-component histidine kinase containing an N-terminal transmembrane domain (TMD), followed by a dimerization histidine phosphotransfer (DHp) domain, a catalytic ATP-binding (CA) domain, and a C-terminal receiver (Rec) domain^2,12^. At low cell densities, when the CAI-1 concentration is below the level of detection, the CqsS CA domain initiates the phosphorylation of an invariant histidine residue (H1) on the DHp domain. Phosphate is next transferred to a conserved aspartate residue (D1) on the CqsS Rec domain. A histidine-containing phosphotransfer (HPt) protein called LuxU subsequently receives the phosphate group from the CqsS D1 residue, again on an invariant histidine residue (H2), and delivers it to an aspartate (D2) on the response regulator called LuxO^13–15^.

Phosphorylated LuxO (LuxO-P) activates the expression of genes encoding quorum regulatory small RNAs (Qrr sRNAs)^16^. The Qrr sRNAs, working in conjunction with the Hfq chaperone, promote the production of the low cell density master transcriptional regulator called AphA and repress the production of the high cell density master transcriptional regulator called LuxR or HapR^16–19^. Under this condition, vibrios behave as individuals (Extended Data Fig. 1a).

At high cell density, accumulated CAI-1 binds to the CqsS TMD, inhibiting its kinase activity^20^. Thus, the CqsS phosphatase activity dominates, resulting in dephosphorylation of LuxO and the termination of Qrr sRNA production^12^. Consequently, AphA production terminates, and the production of LuxR/HapR is derepressed. Under this condition, vibrios undertake group behaviors (Extended Data Fig. 1a).

Despite extensive genetic investigation into CqsS that has identified key residues for signal transduction, the molecular basis underlying catalysis and phospho-relay remain undefined due to a lack of structural information. Here, we determine the cryo-EM structures of full-length *Vibrio harveyi* CqsS in its apo- and ATPγS-bound states. We demonstrate that CqsS is a dimer and autophosphorylation of the H1 residue occurs in *cis* while phosphoryl transfer from CqsS H1 to D1 residues occurs in *trans*. By co-expressing CqsS and the CAI-1 synthase CqsA with the objective of capturing the CAI-1-bound state, we unexpectedly discover a tight interaction between the intracellular domains of CqsS and CqsA, resulting in the formation of a 2:4 complex. In this complex, the H1 residues are positioned away from the CqsS catalytic cavity, a structural observation that supports our discovery of an inhibitory effect of CqsA on CqsS kinase activity. Cryo-EM analysis, focused on the TMD of CqsS in this complex, reveals the presence of two CAI-1 molecules within the TMD, as expected. CAI-1 binding induces obvious movement of transmembrane helix 6, disrupting its extension into the cytosol. This rearrangement provides a possible mechanism for how CAI-1 binding to CqsS in the periplasm is connected to changes in CqsS enzyme activity in the cytoplasm. Together, our findings reveal the molecular mechanism underlying CqsS-CAI-1-mediated QS signal transduction.

## Results

### Structural determination of *Vibrio harveyi* CqsS

The *cqsS* gene was amplified from *V. harveyi* N8T11^21^. The *cqsS* gene and derivatives described throughout this work were introduced into *Escherichia coli* BL21, and the full-length wildtype (WT) and mutant CqsS proteins were purified (see Methods and Extended Data Fig. 1b for WT CqsS). To evaluate CqsS catalytic activity, we assayed autophosphorylation and phospho-relay to LuxU following a reported protocol^12^. WT CqsS efficiently catalyzed both reactions, whereas CqsS H190A, which lacks the crucial H1 residue, exhibited a complete loss of catalytic activity in both assays (Fig. 1a,b). CqsS D613A, which lacks the D1 residue, retained autophosphorylation ability but could not transfer the phosphate to LuxU (Fig. 1a,b). These data align well with previous reports^12^ and confirm the functionalities of our purified proteins.

**Fig. 1.**
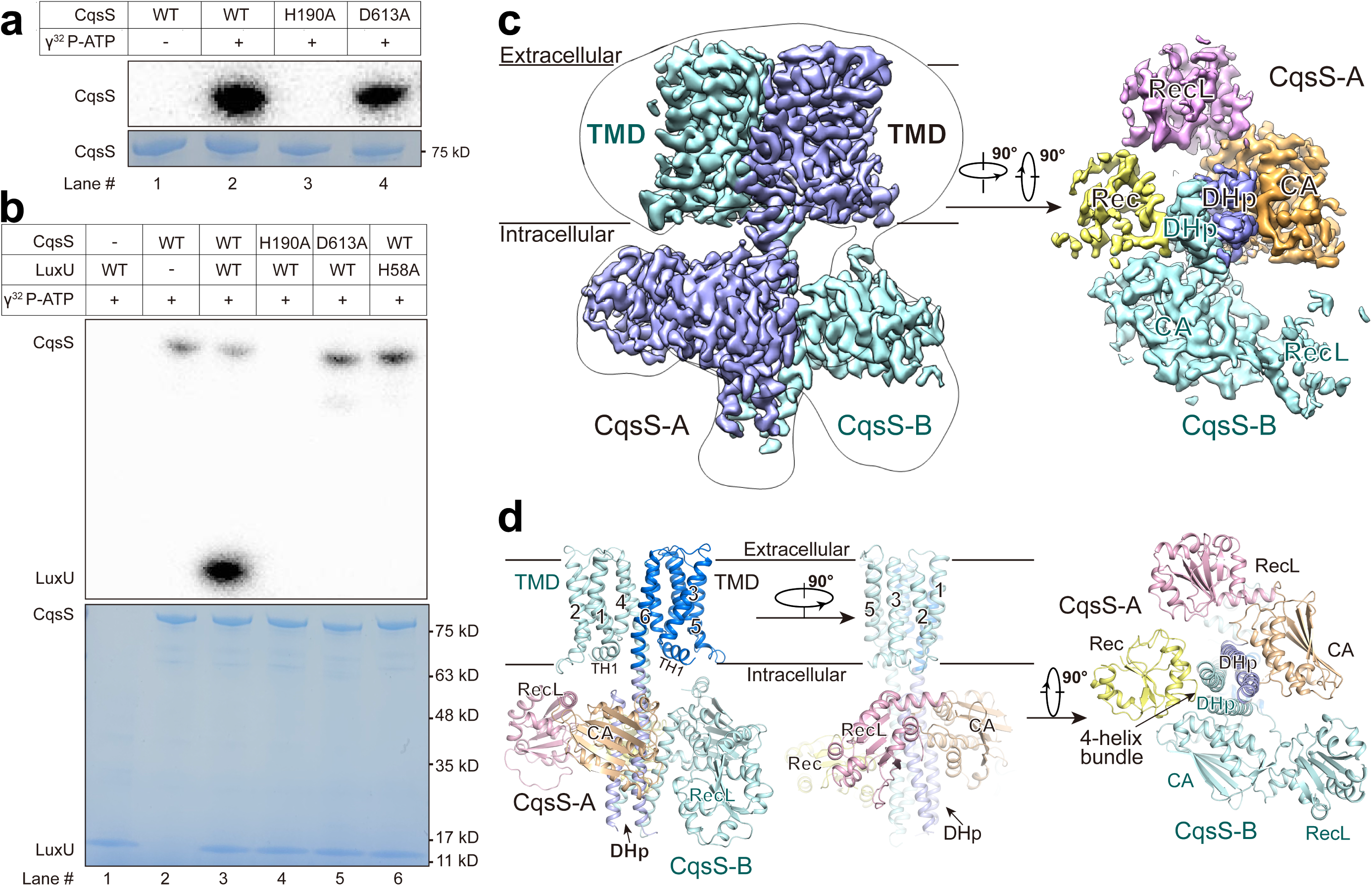
Structural determination of *V. harveyi* CqsS. **a**, Autophosphorylation of CqsS was assessed with the designated purified CqsS proteins. Purified WT and variant CqsS proteins were incubated with radioactive ATP (γ^32^P-ATP) in phosphorylation buffer, followed by autoradiography (top, gray) and Coomassie blue staining (bottom, blue). **b**, Phospho-relay to LuxU was assessed for the designated purified CqsS proteins. Purified WT and variant CqsS proteins were incubated with radioactive ATP (γ^32^P-ATP) and purified LuxU in phosphorylation buffer, followed by autoradiography (top, gray) and Coomassie blue staining (bottom, blue). See Methods for details of the phosphorylation assays. Experiments were repeated more than three times with similar results. **c**, Cryo-EM map of the final reconstructed CqsS_S1_. *Left*: A membrane view to show the dimeric organization, with two protomers colored light blue (CqsS-A) and pale cyan (CqsS-B). A 16 Å low-pass filtered map is shown in transparency to indicate the transmembrane region. *Right*: An intracellular view displays the organization of the cytosolic domains. **d**, Two membrane views (left and middle) and a cytosolic view (right) are presented to show the CqsS_S1_ structure. The CqsS-A protomer is domain-colored with marine blue, light blue, wheat, light pink, and pale yellow for the transmembrane (TMD), dimerization and histidine phosphotransfer (DHp), catalytic and ATP binding (CA), receiver-like (RecL) and receiver (Rec) domains, respectively. The CqsS-B protomer is colored pale cyan. These domains are labeled according to their affiliation with CqsS-A (black) and CqsS-B (dark cyan), respectively, a convention that applies to all figures unless otherwise specified. Each TMD consists of an amino (N)-terminal transverse helix (TH1) and six transmembrane helices (denoted 1-6). The two DHp domains constitute a 4-helix bundle at the dimer interface. All structural images were prepared with PyMOL.

To define the CqsS catalytic mechanism, we employed cryo-EM analysis on purified proteins, which yielded homogenous particles and promising 2D averages (Extended Data Fig. 1c). Following data collection and processing, we obtained two EM maps at resolutions of 3.24 Å (CqsS_S1_, State 1) and 3.28 Å (CqsS_S2_, State 2) with 124,184 and 137,177 particles, respectively (Extended Data Fig. 1d,2a and Supplementary Table 1).

To facilitate model building, we predicted the CqsS structure with AlphaFold3 via the online server (https://alphafoldserver.com/)^22^. The predicted structure was docked into the CqsS_S1_ map. After manual adjustments, the CqsS_S1_ structure was built, which served as the initial model to build the CqsS_S2_ structure. In total, 1,149 and 1,030 residues were assigned to the CqsS_S1_ and CqsS_S2_ structures, respectively (Extended Data Fig. 2a,b and Supplementary Table 1). Given that more residues were assigned to the CqsS_S1_ structure than to CqsS_S2_, unless otherwise noted, our structural analyses focused on the CqsS_S1_ structure.

Our CqsS_S1_ structure shows that CqsS is a dimer, which is consistent with previous reports^1^ (Fig. 1c). Again, consistent with this earlier work, CqsS comprises an N-terminal TMD with 6 transmembrane helices (denoted TM1-6), a DHp domain, a CA domain, and a C-terminal Rec domain (Fig. 1d and Extended Data Fig. 2b-d). Our structures revealed a new cytosolic domain situated between the CA and Rec domains.

We named this new domain, which has a fold similar to that of the Rec domain, the Rec-like (RecL) domain (Fig. 1c,d and Extended Data Fig. 2d-g). In the CqsS dimer, each DHp domain contributes two α-helices to form a 4-helix bundle at the dimer interface (Fig. 1d). The overall structure of the CqsS dimer is asymmetric. Although the transmembrane regions of the two protomers are nearly identical, a 30-degree rotation exists between their DHp domains, resulting in corresponding rotations of the CA and RecL domains (Extended Data Fig. 3a). The RecL domains maintain similar orientations relative to the CA domains in both protomers; however, the Rec domain could only be observed in one protomer (Fig. 1c,d and Extended Data Fig. 3b). For the remainder of this work, to distinguish the two protomers, we designate the protomer with the resolved Rec domain CqsS-A and the other protomer CqsS-B (Fig. 1c,d and Extended Data Fig. 3a,b). Regarding the CqsS_S2_ structure, the Rec domain of CqsS-A is not visible, and the CA and RecL domains in CqsS-B rotate as a rigid body relative to their positions in CqsS_S1_. All other regions are nearly identical between the two structures (Extended Data Fig. 3c,d).

### ATPγS-bound states of CqsS

To define the CqsS autophosphorylation mechanism, we incubated purified CqsS protein with the ATP analog ATPγS to capture the ATPγS-bound state. We obtained two EM maps with resolutions of 3.48 Å and 3.46 Å, designated CqsS-ATPγS_S1_ and CqsS-ATPγS_S2_, respectively (Extended Data Fig. 4 and Supplementary Table 1). After verifying the positions of the known CqsS atoms in the EM maps, we detected additional densities in the ATP binding site of the CA domain in both maps (Fig. 2a and Extended Data Fig. 5a). These densities perfectly accommodate ATPγS molecules (Fig. 2b and Extended Data Fig. 5b), resulting in two distinct structural models, CqsS-ATPγS_S1_ and CqsS-ATPγS_S2_ (Supplementary Table 1). In addition to the ATPγS densities, other densities near the ATPγS molecules are apparent, corresponding to the so-called ATP-lid^23^. The ATP-lid loop could be fully assigned to the CqsS-ATPγS_S1_ structure and partially assigned to the CqsS-ATPγS_S2_ structure (Fig. 2a,c and Extended Data Fig. 5). We therefore focused on the CqsS-ATPγS_S1_ structure to analyze ATPγS coordination.

**Fig. 2.**
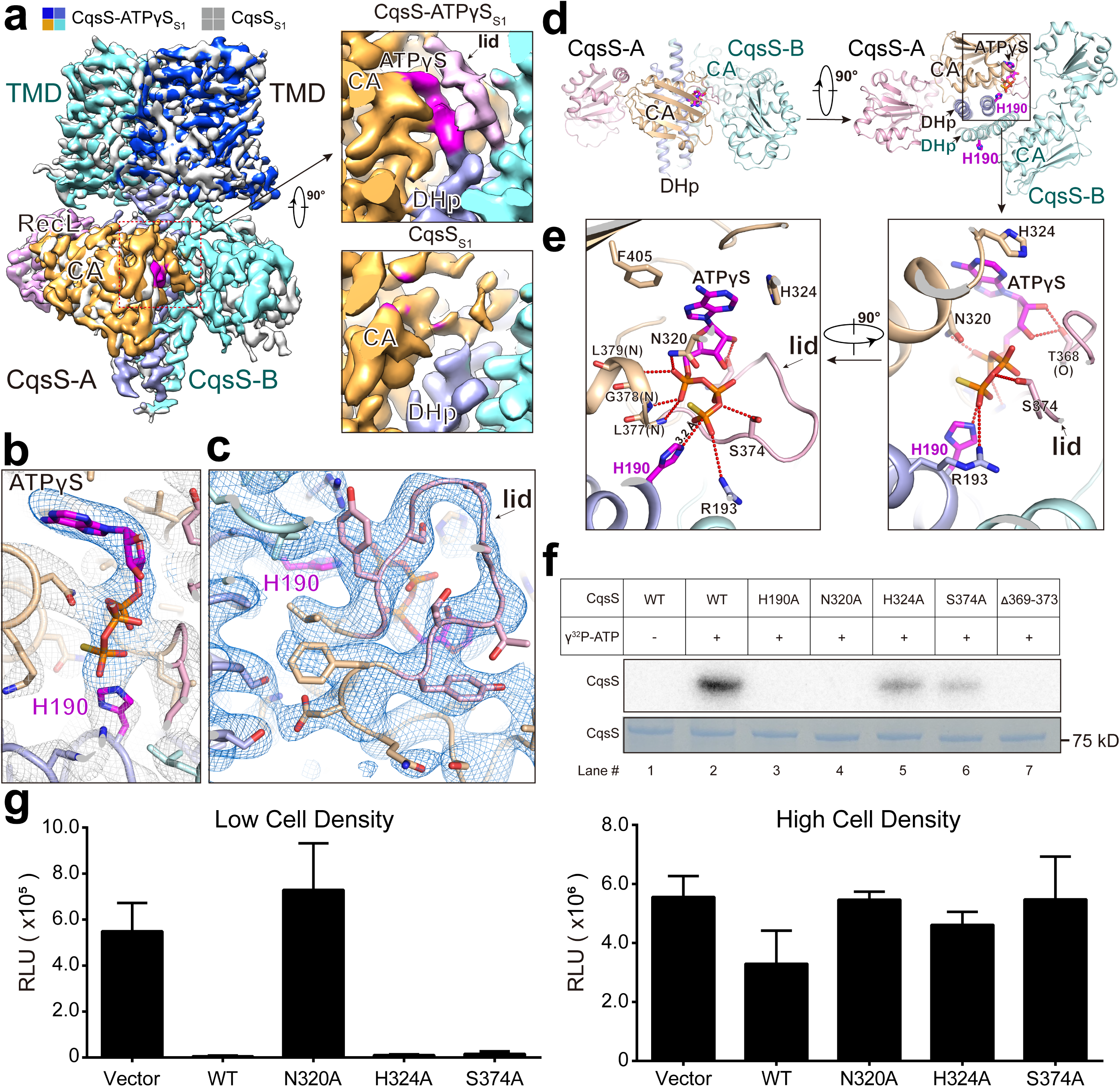
Coordination of ATPγS by the CqsS CA domain. **a**, Overlay of structural maps of CqsS_S1_ and CqsS-ATPγS_S1_. The boxes at the top of the panel indicate that the CqsS-ATPγS_S1_ map is domain-colored (colored box) and the CqsS_S1_ map is presented in gray (gray box). Similar color-coded boxes are also employed to differentiate between maps or structures in other figures. *Insets*: Zoomed in views showing the ATP binding site in the CA domain of the CqsS-A protomer in CqsS-ATPγS_S1_ (top) and CqsS_S1_ (bottom). The densities of ATPγS and the ATP-lid, colored magenta and light pink, respectively, are well resolved in CqsS-ATPγS_S1_. **b**, Structural modeling of an ATPγS molecule into the ATPγS densities shows an excellent fit. The ATPγS densities, contoured at 6σ, are colored marine blue. The CqsS H1 residues (His190) are indicated in magenta in all figures unless otherwise indicated. **c**, Structural modeling of the ATP-lid loop according to the well-resolved densities in the CqsS-ATPγS_S1_ structure. The densities were contoured at 6σ. **d**, The cytoplasmic structure of CqsS-ATPγS_S1_ is presented to show the position of the ATPγS molecule in the cytosolic domain. The CqsS dimer coordinates only one ATPγS molecule in the CA domain of CqsS-A. **e**, Two zoomed in views show the coordination of ATPγS and the role of the ATP-lid loop. Polar interactions are indicated by red dotted lines. **f**, Validation of the essentiality of the designated residues involved in ATPγS coordination in CqsS via assessment of *in vitro* autophosphorylation activity. Purified WT and variant CqsS proteins were incubated with radioactive ATP (γ^32^P-ATP) in phosphorylation buffer, followed by autoradiography (top, gray) and Coomassie blue staining (bottom, blue). The experiments were independently repeated more than three times with similar results. **g**, Validation of the essentiality of the designated residues involved in ATPγS coordination in CqsS via assessment of *in vivo* kinase activity in Δ*cqsS V. harveyi* carrying different CqsS variants. The graphs present the minimum and maximum light outputs in the assay, which occur at low cell density (left) and high cell density (right), respectively. RLU refers to relative light units, which is bioluminescence per OD_600_. Error bars denote standard deviations of the means, *n*=3. See the Methods section for all the assay conditions.

Despite the presence of two protomers in the CqsS-ATPγS_S1_ structure, the ATPγS molecule could only be observed in the CA domain of the CqsS-A protomer (Fig. 2d). In this protomer, the CA domain is located near the H1 residue (His190) on the DHp domain, whereas the CA domain of the CqsS-B protomer is located distal to the H1 residue on its DHp domain (Fig. 2d). This structural observation suggests that histidine phosphorylation occurs in one protomer at a time, which is consistent with previous reports on other histidine kinases^24,25^.

The ATPγS molecule is primarily coordinated within the CA domain through polar interactions. Specifically, the phosphate groups on the ATPγS molecule form hydrogen bonds with the side chains of CqsS-A Ser374 and Asn320, as well as with the main chain amides of CqsS-A residues 377-379. The ATPγS ribose moiety potentially engages in hydrogen bonding with the carbonyl group in the main chain of CqsS-A Thr368. Furthermore, the ATPγS adenosine ring is stabilized by π-interactions with CqsS-A His324 and Phe405 (Fig. 2e). To validate our structural observations, we introduced alanine substitutions into CqsS to disrupt coordination of ATPγS, resulting in the CqsS N320A, CqsS H324A, and CqsS S374A variants. Alanine substitution at the H1 residue (CqsS H190A) was also included as the control. Robust autophosphorylation occurs with WT CqsS following the addition of ATP, whereas the CqsS H190A variant does not undergo autophosphorylation (Fig. 2f, lanes 1-3; Extended Data Fig. 6a). Regarding ATP-coordination, mutation at CqsS N320 completely abolished *in vitro* autophosphorylation, and CqsS H324A and CqsS S374A displayed severely diminished autophosphorylation (Fig. 2f, lanes 4-6; Extended Data Fig. 6a), supporting their essential roles in ATP coordination. We also deleted the ATP-lid (CqsS Δ369-373) to examine its role in autophosphorylation. This CqsS variant also lost the ability to catalyze autophosphorylation, showing that the ATP-lid is indispensable (Fig. 2f, lane 7; Extended Data Fig. 6a).

To probe the *in vivo* activities of our CqsS mutants, we introduced them into Δ*cqsS V. harveyi* and assessed their effects on bioluminescence emission, the hallmark QS-controlled trait. Specifically, in WT *V. harveyi*, at low cell density, when endogenously produced CAI-1 levels are low, CqsS kinase activity represses light production (Fig. 2g, Low Cell Density). At high cell density, in response to CAI-1 binding, CqsS kinase activity is inhibited and CqsS phosphatase activity promotes light production (Fig. 2g, High Cell Density). The left panel of Fig. 2g shows that, at low cell density, when the vector alone is introduced into Δ*cqsS V. harveyi*, high-level light production occurs. This is because with no CqsS kinase activity to suppress bioluminescence, the default state of the system is constitutive light production. Indeed, introduction of WT CqsS represses light production at low cell density. CqsS N320A, which is completely defective in ATP binding/autophosphorylation (Fig. 2f), is also constitutively bright, showing that there is no *in vivo* CqsS kinase activity (Fig. 2g and Extended Data Fig. 6d). By contrast, Δ*cqsS V. harveyi* carrying CqsS H324A and CqsS S374A are dark at low cell density (Fig. 2g and Extended Data Fig. 6d), demonstrating that the residual autophosphorylation/kinase activity shown in Fig. 2f is sufficient for full repression of bioluminescence *in vivo*. The right panel of Fig. 2g shows that WT CqsS and the three mutants discussed here all display phosphatase activity at high cell density.

Fig. 2e shows that the ATP γ-phosphate is located at a distance of 3.2 Å from the εN atom of the H1 (His190) residue in the DHp domain, presumably in proximity to mimic nucleophilic attack of the εN atom on the ATP γ-phosphate. The CqsS Arg193 residue in the DHp domain also engages in electrostatic interactions with the ATP γ-phosphate group, further stabilizing the ATP γ-phosphate relative to the H1 residue (Fig. 2e). Analogous to other bacterial histidine kinases, the proton on the H1 residue may be abstracted by a conserved negatively charged residue (in CqsS that is Glu191) to facilitate nucleophilic attack (Extended Data Fig. 5d)^25–27^. Our structure shows that CqsS Glu191 also interacts with Asn316 on the CA domain, the latter presumably aiding in maintaining the optimal relative orientations of these two domains (Extended Data Fig. 5d). In support of these speculations, CqsS E191A and CqsS N316A completely lack the ability to catalyze autophosphorylation (Extended Data Fig. 6b,c) and to repress bioluminescence at low cell densities *in vivo* (Extended Data Fig. 6d,e).

The CqsS-ATPγS_S2_ structure shows that CqsS-A displays a similar structural arrangement to that described above to coordinate the ATPγS molecule, even though only a portion of its ATP-lid could be resolved (Extended Data Fig. 5e). In the CqsS-ATPγS_S1_ structure, the ATP-lid appears to be stabilized by a salt bridge between CqsS Lys371 and CqsS Glu258 from the CA domain in the other subunit (Extended Data Fig. 5e, bottom left). However, this salt bridge is disrupted in the CqsS-ATPγS_S2_ structure, potentially leading to the partially resolved ATP-lid loop (Extended Data Fig. 5e, bottom right; Supplementary Video 1). As described above, the ATP-lid contributes Thr368 and Ser374 residues to the coordination of ATPγS (Fig. 2e and Extended Data Fig. 5e). Therefore, a salt bridge between two CqsS CA domains appears crucial for kinase activity because it maintains the appropriate conformation of the ATP-lid. To validate this hypothesis, we substituted Lys371 with alanine (K371A) in CqsS. The CqsS K371A variant exhibited significantly diminished autophosphorylation (Extended Data Fig. 6b,c), supporting our structural predictions. Nonetheless, and similar to CqsS H324A and CqsS S374A, the residual kinase activity present in CqsS K371A is sufficient to repress *in vivo* bioluminescence (Extended Data Fig. 6d,e).

Comparisons of the ATPγS-bound structures (CqsS-ATPγS_S1_ and CqsS-ATPγS_S2_) with the apo-structures (CqsS_S1_ and CqsS_S2_), show obvious conformational changes in the RecL domain of CqsS-B. This domain appears to move progressively away from the transmembrane domain, from CqsS-ATPγS_S1_ through CqsS_S2_ and CqsS-ATPγS_S2_ to CqsS_S1_ (Extended Data Fig. 5f and Supplementary Video 1). Detailed examination of these structures reveals a modest rotation of the CA domain in CqsS-B. This rotation likely causes the pronounced shift in the RecL domain of CqsS-B. This conformational change may help stabilize the Rec domain of CqsS-A, resulting in the observed Rec domain of CqsS-A in the CqsS_S1_ structure (Extended Data Fig. 5f).

### *Cis*-phosphorylation of the CqsS H1 residue

Autophosphorylation of bacterial histidine kinases can occur in *trans* or in *cis*, depending on the handedness of the DHp domains^28^. Our structures appear to support *cis*-phosphorylation of the H1 residue because the bound ATPγS molecule is oriented toward the H1 (His190) residue of the same subunit in the CqsS-ATPγS_S1_ structure (Fig. 3a).

**Fig. 3.**
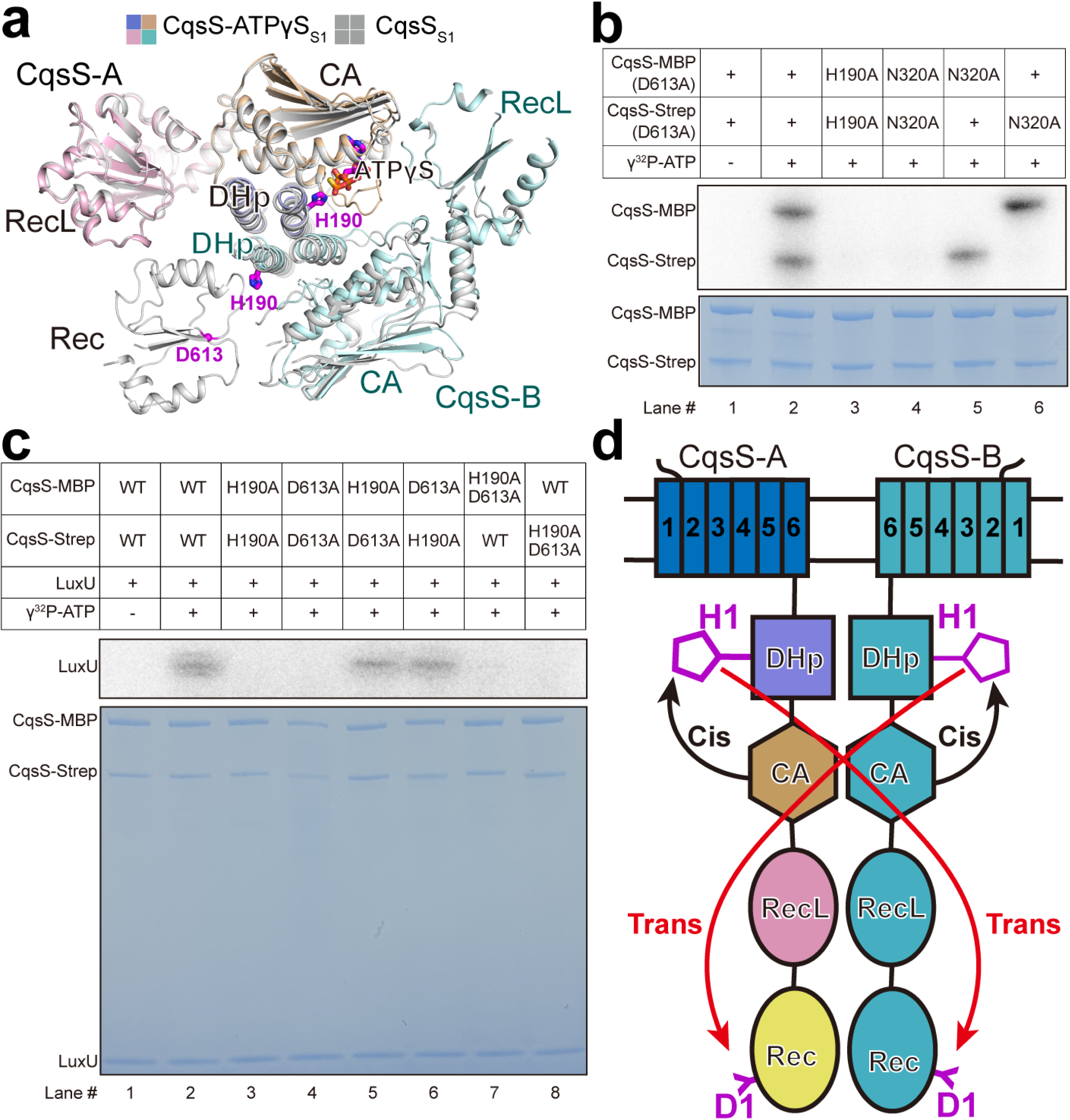
*Cis* phosphorylation of His190 and *trans* phospho-transfer from His190 to Asp613 occur in CqsS. **a**, Structural overlay of the cytoplasmic domains of CqsS_S1_ and CqsS-ATPγS_S1_. In protomer A, ATPγS is coordinated near the H1 residue (His190) of that protomer, indicating *cis* phosphorylation. The Rec domain of protomer A lies adjacent to the H1 residue of the CqsS-B protomer, suggesting *trans* phospho-transfer from CqsS H1 to D1 (Asp613). Both H1 and D1 residues are colored magenta. **b**, Validation of *cis*-phosphorylation of the CqsS H1 residue via assessment of *in vitro* autophosphorylation activity. The CqsS chimeras harboring the designated substitutions were incubated with radioactive ATP (γ^32^P-ATP) in phosphorylation buffer, followed by autoradiography (top, gray) and Coomassie blue staining (bottom, blue). **c**, Functional validation of *trans*-phospho-transfer from CqsS H1 to D1 residues. The CqsS chimeras harboring the designated substitutions were incubated with radioactive ATP (γ^32^P-ATP) and purified LuxU protein in phosphorylation buffer, followed by autoradiography (top, gray) and Coomassie blue staining (bottom, blue). **d**, A proposed model illustrating *cis* phosphorylation of the CqsS H1 residue and *trans*-phospho-transfer to the D1 residue. The TMD segments and cytoplasmic domains are labeled as in Fig. 1. Black and red arrows indicate *cis* and *trans* phospho-transfer, respectively. The experiments in panels **b** and **c** were independently conducted at least three times with similar results. See the Methods section for details on the purification of the CqsS-Strep and CqsS-MBP chimeras and the assay conditions.

To validate this structural observation, we conducted *in vitro* autophosphorylation experiments using chimeric versions of CqsS in which the individual subunits were fused to either Strep or MBP tags. Following tandem affinity purification, CqsS chimeras were confirmed to be homogeneous via size exclusion chromatography (SEC) analysis (Extended Data Fig. 6f). Additionally, we substituted Asp613 (D1) with alanine in both subunits to prevent phospho-relay from H1 to D1. The resulting CqsS chimera effectively catalyzed autophosphorylation (Fig. 3b, lanes 1-2). When H1 residues were substituted with alanine (H190A) in both subunits, no phosphorylation was detected in either subunit, as expected (Fig. 3b, lane 3). A similar phenomenon was observed when the kinase inactivating N320A mutation was introduced into the CA domains of both subunits (Fig. 3b, lane 4). To differentiate between the *cis*- or *trans*-phosphorylation mechanism of the H1 residue, the N320A substitution was introduced into only one subunit. Consistent with our structural observations, this mutation abolished phosphorylation only on the *cis* subunit, but not on the *trans* subunit (Fig. 3b, lanes 5-6), thereby confirming a *cis*-phosphorylation mechanism on the H1 residue.

### *Trans*-phospho-relay from CqsS H1 to D1 residue

Following phosphorylation at the H1 residue, the phosphate group is transferred to the D1 (Asp613 in CqsS) residue located on the Rec domain in bifunctional two-component proteins like CqsS. In the CqsS_S1_ structure, the Rec domain is positioned adjacent to the H1 residue from the other subunit (Fig. 3a), suggesting a *trans*-phospho-transfer reaction from CqsS H1 to D1.

To confirm our structural observations, we performed the *in vitro* phosphorylation assay, again exploiting CqsS chimeras for the analysis. Owing to the challenge of distinguishing between H1 and D1 phosphorylation within the same subunit, we measured phosphorylation of LuxU, the downstream phosphate acceptor, as an indicator of successful phospho-transfer from CqsS H1 to D1. When both CqsS H1 residues were altered to alanine, CqsS lost the ability to catalyze LuxU phosphorylation, confirming the critical role of H1 in the initial autophosphorylation step (Fig. 3c, lane 3). A similar loss of function occurred when both CqsS D1 residues were mutated, underscoring their importance in the phospho-transfer steps (Fig. 3c, lane 4). To distinguish between *cis* and *trans* mechanisms of phospho-transfer from CqsS H1 to D1, we constructed CqsS chimeras in which we substituted a single H1 residue and a single D1 residue with alanine, either within the same subunit or the mutations were made on separate subunits. WT levels of LuxU phosphorylation occurred when the CqsS H1 and D1 mutations were in different subunits (Fig. 3c, lanes 5-6; Extended Data Fig. 6g). By contrast, when the mutations were present within the same subunit, CqsS exhibited no phospho-transfer to LuxU (Fig. 3c, lanes 7-8). This pattern confirms that phospho-transfer from CqsS H1 to D1 occurs in *trans* between different subunits.

Collectively, our structural observations and companion biochemical characterization demonstrate that CqsS autophosphorylation initiation occurs in *cis* at the H1 residue, whereas phospho-transfer from CqsS H1 to the D1 residue occurs in *trans* (Fig. 3d).

### Direct interaction occurs between CqsS and the CAI-1 synthase CqsA

Garnering information concerning the mechanism used by CqsS to detect CAI-1 requires the structure of CqsS bound to CAI-1. Fortuitously, when the *cqsA* gene encoding the *V. harveyi* CAI-1 synthase CqsA is introduced into *E. coli*, it produces CAI-1^10^. We therefore co-expressed *cqsS* and *cqsA* in *E. coli*, anticipating purification of CqsS in the CAI-1-bound state. To our surprise, we identified an additional protein that co-purified with CqsS. The co-purifying protein had a molecular weight of approximately 50 kDa, closely matching that of CqsA. Moreover, this protein co-migrated with CqsS during SEC analysis, indicating a tight interaction (Extended Data Fig. 7a). Western blot analysis showed that the CqsS band was immunostained with both anti-Strep and anti-FLAG antibodies, consistent with the fact that both Strep and FLAG tags were fused in tandem to the C-terminus of CqsS. By contrast, on Western blots, the additional band could only be detected with anti-FLAG antibody, confirming this protein is CqsA, which carried a FLAG tag at the N-terminus (Extended Data Fig. 7a). These observations suggest that a direct interaction occurs between the CqsS receptor and the CAI-1 synthase CqsA.

We used an *in vitro* pull-down assay to validate the interaction between CqsS and CqsA. Strep-tagged, purified CqsS was incubated with independently purified CqsA, followed by affinity purification using Strep-Tactin resin. The presence of CqsA in the eluate confirmed the direct interaction between CqsS and CqsA (Fig. 4a). Critically, CqsA K240A, which corresponds to the catalytically-inactive *V. cholerae* CqsA K236A mutant protein^29^, demonstrated an ability similar to WT CqsA to interact with CqsS (Extended Data Fig. 7b). Furthermore, interaction with CqsA clearly inhibits CqsS autophosphorylation and subsequent phospho-relay to LuxU. CqsA catalytic activity is also not required as the CqsA K240A variant displays the same inhibitory ability (Fig. 4b, c).

**Fig. 4.**
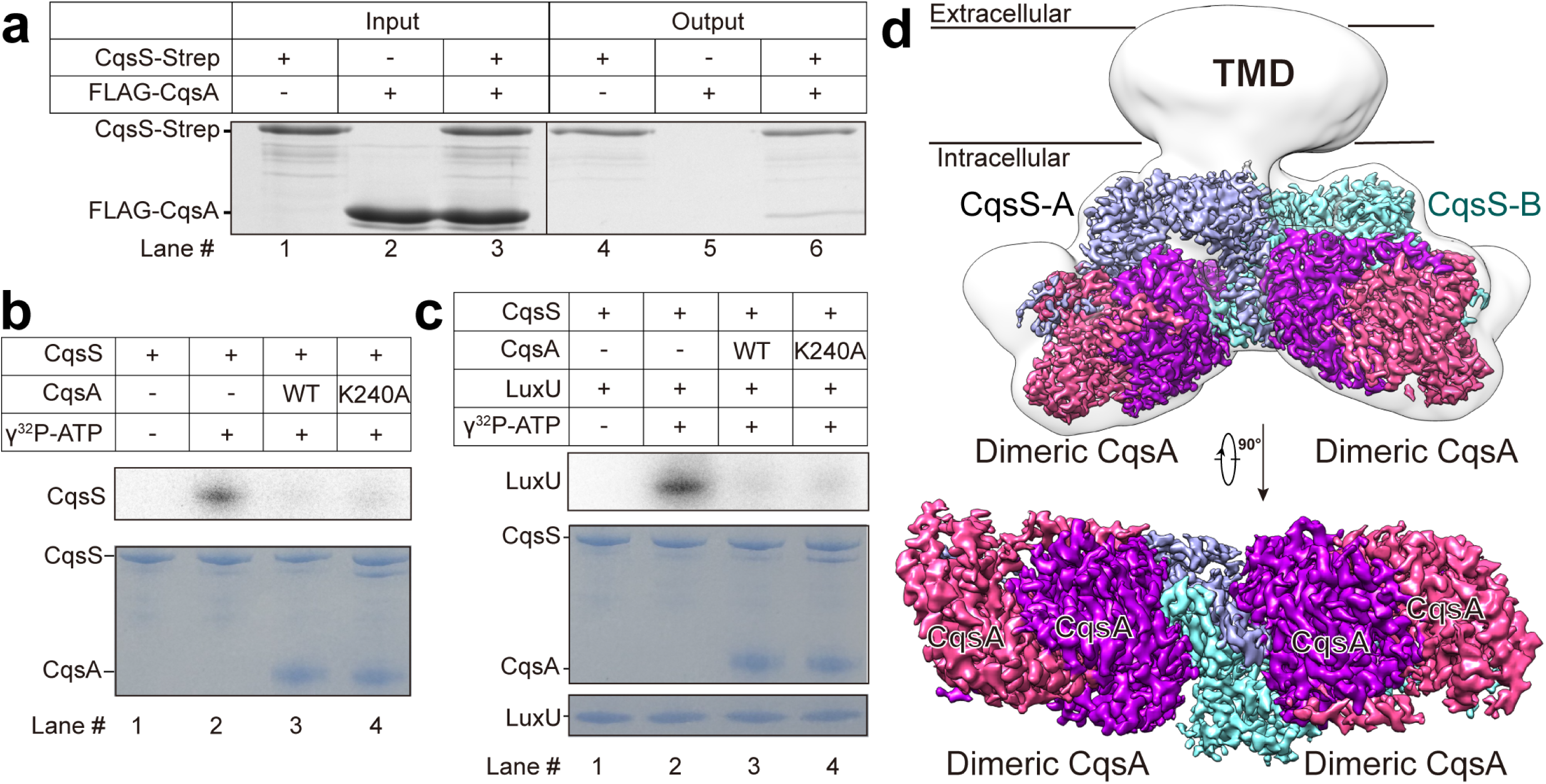
A direct interaction occurs between CqsS and the CAI-1 synthase CqsA. **a**, Direct interaction between CqsS and CqsA was validated via an *in vitro* pull-down assay employing a Strep tag on CqsS. The input (lanes 1-3) and output (lanes 4-6) of the pull-down assay were subjected to SDS-PAGE analysis, followed by Coomassie blue staining. See Methods for details. **b**, CqsA inhibits CqsS autophosphorylation independently of its catalytic activity. Purified CqsS protein was incubated with radioactive ATP (γ^32^P-ATP) and either WT or the catalytically-inactive CqsA K240A protein in phosphorylation buffer, followed by autoradiography (top, gray) and Coomassie blue staining (bottom, blue). **c**, CqsA inhibits phospho-relay from CqsS to LuxU independently of its catalytic activity. WT or the catalytically-inactive CqsA K240A protein was incubated with purified CqsS protein, radioactive ATP (γ^32^P-ATP), and LuxU in phosphorylation buffer, followed by autoradiography (top, gray) and Coomassie blue staining (bottom, blue). **d**, Membrane (top) and cytosolic (bottom) views showing the EM map of the CqsS-CqsA complex. A 16 Å low-pass filtered map is shown in transparency under a low threshold to indicate the TMD. Because of the high flexibility between the cytosolic and transmembrane domains, the TMD is invisible at a high threshold. The two CqsA subunits in the CqsA dimer are colored dark purple and magenta, and the CqsS-A and CqsS-B protomers are colored light blue and pale cyan.

### Structural determination of the CqsS-CqsA and CqsS-CAI-1 complexes

To probe the interaction between CqsS and CqsA and to capture the structure of CqsS in the CAI-1-bound state, we subjected the purified complex of CqsS and CqsA to cryo-EM analysis, resulting in homogenous particles suitable for further cryo-EM structure determination (Extended Data Fig. 7c). We collected a large dataset, and after 2D classification, we identified two distinct sets of 2D averages: one displaying a clear intracellular domain (ICD) but a blurred TMD and another with a clear TMD but a blurred ICD (Extended Data Fig. 7d). This observation suggests significant flexibility between the ICD and TMD. These two types of 2D averages were chosen as templates for particle selection, with the aim of determining separate cryo-EM maps for the ICD and TMD. Indeed, we successfully obtained two EM maps at resolutions of 2.87 Å and 3.38 Å for the ICD and TMD, respectively (Extended Data Fig. 7e-f,8 and Supplementary Table 1). The cytosolic domains of two CqsS protomers and two CqsA dimers were subsequently built into the ICD map, resulting in a 2:4 complex structure designated CqsS-CqsA (Fig. 4d). The CAI-1 molecules were resolved in the TMD map, resulting in a CAI-1-bound structure for CqsS, which we designate CqsS-CAI-1. Below, we first focus on the CqsS-CqsA map to analyze the interaction between CqsS and CqsA, and after that, we discuss the EM map of the TMD and what it reveals about CAI-1 binding.

### Analysis of the CqsS-CqsA interaction

In the CqsS-CqsA structure, each CqsS protomer interacts with a CqsA dimer to generate a 1:2 subcomplex. Two subcomplexes are connected via the DHp domains to generate the 2:4 complex (Fig. 5a). In contrast to the distinct conformations adopted by the two protomers in the CqsS_S1_ structure (Extended Data Fig. 3a), in the CqsS-CqsA structure, the two subcomplexes are nearly identical, with an RMSD of 0.3 Å for 1,114 Ca atoms (Extended Data Fig. 9a). Notably, compared to the CqsS_S1_ structure, the Rec domain undergoes a significant rearrangement relative to the CA and RecL domains in the CqsS-CqsA structure. This rearrangement alters the distance between the D1 residue and the *trans* H1 residue from 18 Å to 85 Å (Extended Data Fig. 9b), indicating disruption of *trans* phospho-transfer from H1 to D1. Moreover, the DHp domain also exhibits a movement of approximately 10 Å relative to the CA domain when comparing the CqsS-CqsA and CqsS-ATPγS_S1_ structures. This movement also increases the distance between the γ-phosphate of ATPγS and the εN atom of the H1, extending it from 3.2 Å to 10 Å, which provides a structural explanation for the inhibitory effect of CqsA on CqsS kinase activity (Fig. 4b,c and Extended Data Fig. 9b). We note that the DHp domain is better resolved in the CqsS-CqsA structure than in the CqsS_S1_ structure. In the CqsS-CqsA structure, the loops connecting α1 and α2 helices of the DHp domain are well-defined unlike in the CqsS_S1_ structure (Extended Data Fig. 9c).

**Fig. 5.**
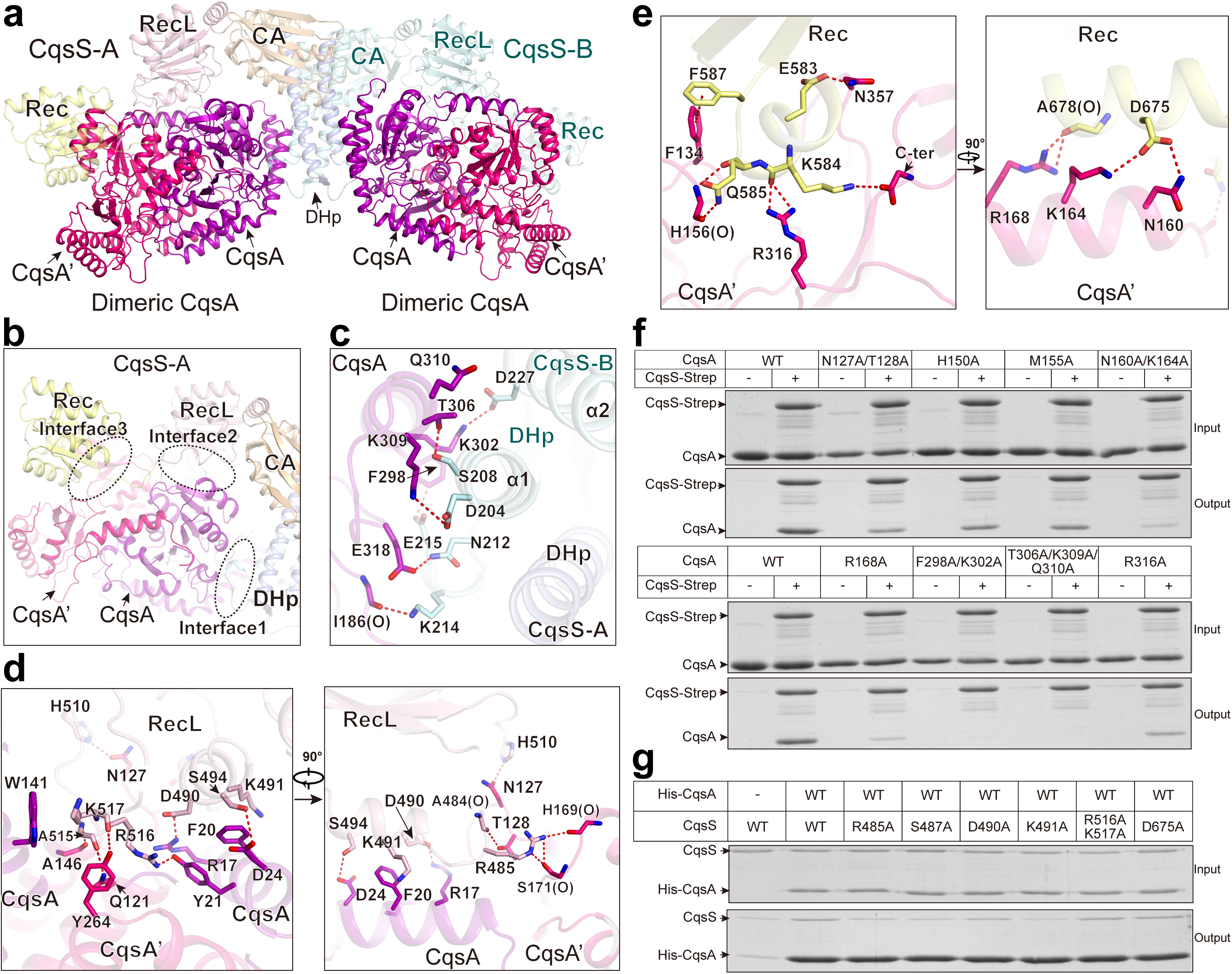
CqsS and CqsA form a complex via three interaction interfaces. **a**, The cytosolic domain of each CqsS protomer binds to one dimer of CqsA, resulting in a 2:4 complex. The two CqsA protomers in each CqsA dimer are labeled CqsA and CqsA’ and are colored dark purple and magenta, respectively. **b**, Dimeric CqsA interacts with the CqsS DHp, RecL, and Rec domains using three interfaces, labeled Interface 1, 2, and 3. **c**-**e**, The interaction details between dimeric CqsA and the DHp (**c**), RecL (**d**), and Rec (**e**) domains. Each dimeric CqsA protein interacts with the RecL and Rec domains from the same CqsS subunit and with the DHp domain of the other subunit. The “O” character in the labels of particular residues indicates that the main chain carboxyl groups of these residues participate in hydrogen bond formation. **f**, Biochemical validation of essential residues on CqsA involved in the CqsS-CqsA interaction via *in vitro* pull-down assays employing Strep tagged CqsS. **g**, Biochemical validation of critical residues on CqsS involved in the CqsS-CqsA interaction via *in vitro* pull-down assays employing 10xHis tagged CqsA.

In the CqsS-CqsA structure, each CqsA dimer engages in three distinct interfaces, which we name interfaces 1, 2, and 3, enabling it to interact with the DHp, RecL, and Rec domains, respectively (Fig. 5b, ovals). For clarity, we designate the two subunits in the CqsA dimer CqsA (which participates in interfaces 1 and 2) and CqsA’ (which engages in interfaces 2 and 3) (Fig, 5b). At interface 1, the CqsA subunit interacts with the CqsS DHp domain from a *trans* subunit via five polar interactions, including CqsS Lys214-CqsA Ile186, CqsS Asp227-CqsA Lys302, CqsS Ser208-CqsA Thr306, CqsS Asp204-CqsA Lys309, and CqsS Asn212-CqsA Glu318. Additionally, Phe298 on CqsA likely forms a hydrogen bond with Glu215 on the *trans* CqsS DHp domain (Fig. 5c). At interface 2, the CqsS RecL domain employs several charged residues, Arg485, Asp490, Lys491, Arg516 and Lys517, to interact with polar residues from both subunits of the CqsA dimer (Fig. 5d). Interface 3 is characterized by multiple hydrogen bonds between CqsS and CqsA’, including CqsS Glu583-CqsA’ Asn357, CqsS Gln585-CqsA’ His156, CqsS Gln585-CqsA’ Arg316, CqsS Asp675-CqsA’ Asn160 and CqsS Ala678-CqsA’ Arg168. Additionally, this interface features several salt bridges, including CqsS Asp675-CqsA’ Lys164 and CqsS Lys584 with the C-terminal carboxyl group of CqsA’, and a π‒π interaction between CqsS Phe587 and CqsA’ Phe134 (Fig. 5e).

To validate the importance of these interfaces, single, double, and triple mutations were introduced into CqsS and CqsA to disrupt them, and the consequences for CqsS-CqsA complex formation were assessed using *in vitro* pull-down assays. Most mutations impaired (CqsA N127A/T128A, H150A, M155A, N160A/K164A, R168A, R316A and CqsS R485A, S487A, K491A) or eliminated (CqsA F298A/K302A, T306A/K309A/Q310A) the interaction between CqsS and CqsA in our assay, confirming the essential roles of these residues in forming the CqsS-CqsA interfaces (Fig. 5f,g and Extended Data Fig. 9d). By contrast, the CqsS mutations D490A, R516A/K517A, and D675A had negligible effects on the interaction between CqsS and CqsA (Fig. 5g and Extended Data Fig. 9d).

### CAI-1 coordination by the CqsS-TMD

As mentioned above, CAI-1 was present in the EM map of the TMD of the purified complex following co-expression of CqsS and CqsA, consistent with its hydrophobic nature and previous genetic analyses^12,20^. Our examination identified two CAI-1-shaped densities in the TMD at the interface between the two CqsS protomers (Fig. 6a; 6b, left). We therefore modeled two CAI-1 molecules into these extra densities (Fig. 6c), resulting in a structure that, above, we designated CqsS-CAI-1. Additional densities are also present at similar positions in the apo-structure (CqsS_S1_), but their shapes differ from those of the CAI-1 molecule. Lipid molecules, such as diacylglycerol, fit more appropriately into these additional densities, suggesting that other hydrophobic compounds may occupy these sites within the CqsS_S1_ structure (Fig. 6b, right and Extended Data Fig. 10a,b).

**Fig. 6.**
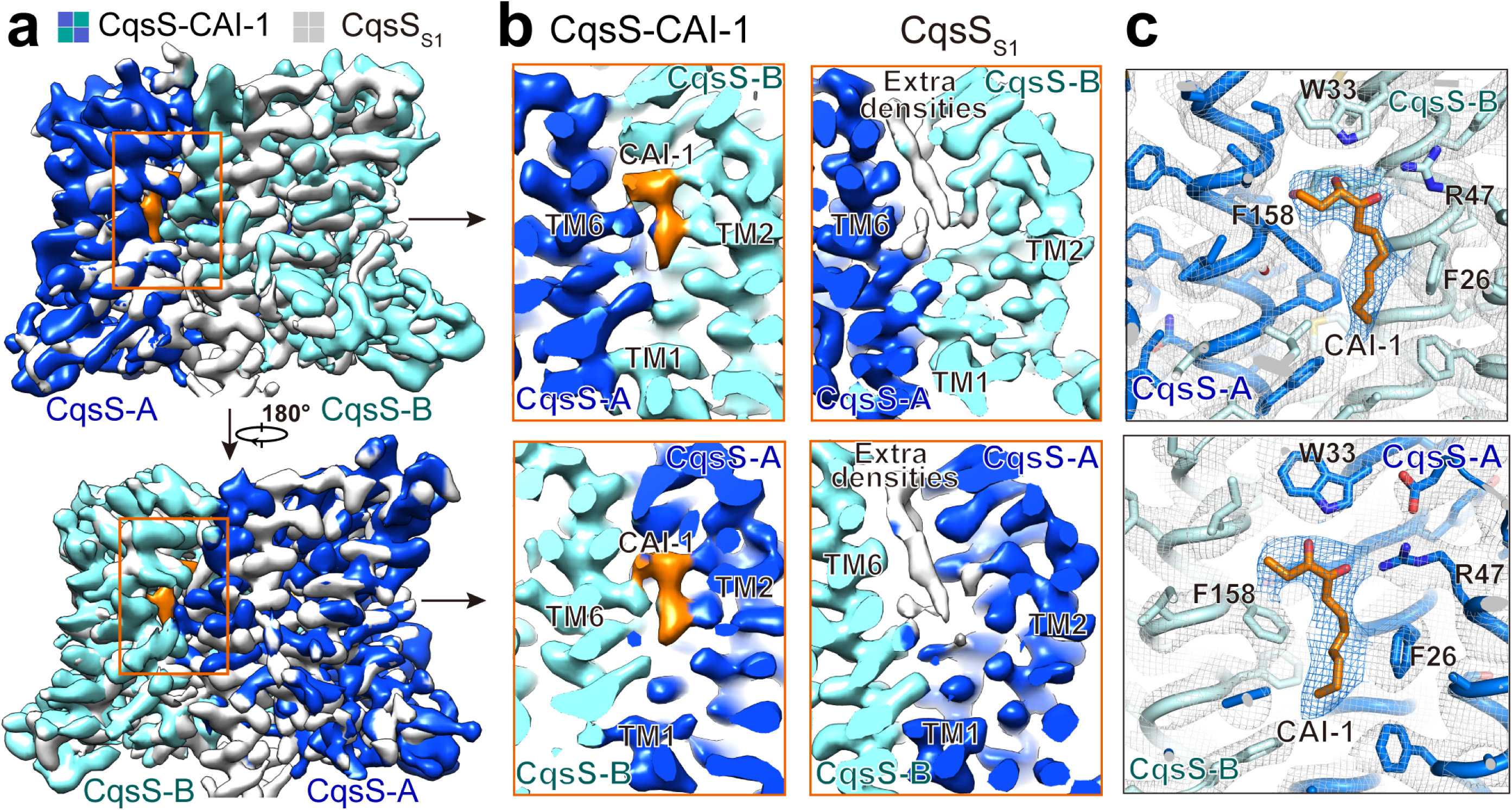
CAI-1 densities are observed in the transmembrane region of the CqsS-CAI-1 structure. **a**, Overlay of the TMD maps between CqsS-CAI-1 and CqsS_S1_. Two CAI-1 molecules (orange) are located in the CqsS-CAI-1 structure, with each molecule primarily coordinated by CqsS-A (top) and CqsS-B (bottom), respectively. **b**, EM densities for two CAI-1-like moieties exist at the dimer interface within the TMD of the CqsS-CAI-1 structure (left), and additional densities that do not match those assigned to CAI-1 are also present at similar sites in CqsS_S1_ (right). Both maps are low-pass filtered to 3.4 Å and set at an 8σ cutoff for comparison. CAI-1 molecules are colored orange. **c**, Structural modeling of CAI-1 molecules into CAI-1-like densities within the CqsS-CAI-1 structure. The CAI-1 densities, contoured at 7σ, are colored in marine blue, and CAI-1 molecules are shown as orange sticks.

In the CqsS-CAI-1 structure, the CAI-1 molecules are located on the extracellular side of the TMD. The two CAI-1 molecules are symmetrically coordinated by TM1-4 from one protomer and TM6 from the other protomer (Fig. 7a). Thus, TM1-4 appear to provide the primary CAI-1 binding site. The polar CAI-1 headgroup forms hydrogen bonds with CqsS TM1 Trp33, TM2 Arg47, and TM4 Ser103, and the nonpolar CAI-1 fatty acyl tail is stabilized by hydrophobic residues from TM1 (Phe26 and Tyr29) and TM3 (Leu83, Phe86, and Phe87). TM6 from the adjacent CqsS protomer also contributes several hydrophobic residues (Phe158, Phe162, and Phe166) to stabilize the hydrophobic tail (Fig. 7b). In support of our structural observations, a previous functional study suggested that residues Phe91, Ser107, and Phe162 in *V. cholerae* CqsS (designated *Vc*CqsS), corresponding to TM3 Phe87, TM4 Ser103, and TM6 Phe158, respectively, in *V. harveyi* CqsS, are essential for CAI-1 recognition^12,20^. Moreover, *Vc*CqsS Ser107 was reported to determine the preference for the moiety on the CAI-1 C3 atom^20^. In our *V. harveyi* CqsS-CAI-1 structure, CqsS Ser103, which as mentioned, corresponds to *Vc*CqsS Ser107, forms a hydrogen bond with the hydroxyl group on C3 of CAI-1, providing a structural explanation for this earlier biochemical finding (Fig. 7b).

**Fig. 7.**
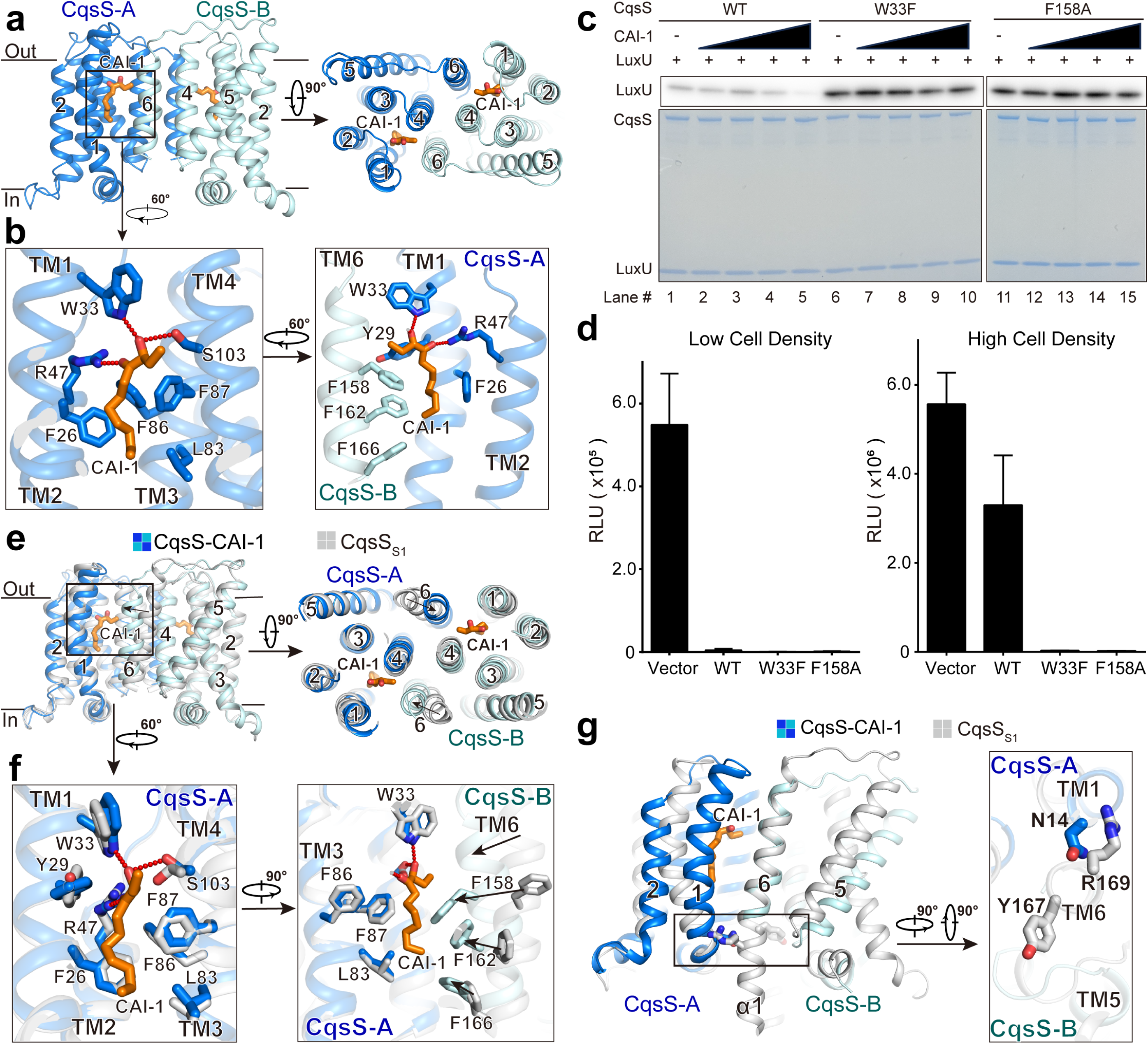
The CqsS CAI-1 sensing mechanism. **a**, A membrane view and an extracellular view of the CqsS TMD highlight the position of CAI-1 within the TMD. Two CAI-1 molecules are located in the TMD that resides in the outer leaflet of the cytoplasmic membrane, each surrounded by TMs 1-4 from one CqsS protomer and TM6 from another. **b**, Two zoomed in views show the details of CAI-1 coordination in the CqsS-A protomer. Hydrogen-bonds are indicated by red dotted lines. **c**, *In vitro* validation of essential residues involved in CAI-1 coordination to CqsS via a CqsS phospho-relay assay. The designated purified CqsS proteins were incubated with radioactive ATP (γ^32^P-ATP), increasing amounts of cell-free culture medium containing CAI-1 (ranging from 0.031% to 2.0% (v/v) of the original cell-free culture fluids), and purified LuxU in phosphorylation buffer, followed by autoradiography (top, gray) and Coomassie blue staining (bottom, blue). **d**, *In vivo* validation of essential residues involved in CqsS binding to CAI-1 and consequent inhibition of CqsS kinase activity in Δ*cqsS V. harveyi* carrying WT CqsS and the designated CqsS variants. The graphs present the minimum and maximum light output in the assay, which occur at low cell density (left) and high cell density (right), respectively. RLU refers to relative light units which is bioluminescence per OD_600_. Error bars denote standard deviations of the means, *n*=3. See the Methods section for all assay conditions. **e**, A membrane view and an extracellular view show overlays of the TMDs between CqsS-CAI-1 and CqsS_S1_. Note especially the rearrangement of TM6. **f**, Two zoomed-in views enable comparison of the CAI-1 binding sites between the apo- and CAI-1-bound structures. The CqsS-CAI-1 and CqsS_S1_ structures are superimposed relative to the TMD of the CqsS-A protomer. **g**, Structural overlay of CqsS-CAI-1 and CqsS_S1_ relative to TM6 in the CqsS-B protomer. Steric clashes between TM1 and TM6 from another subunit, presumably driven by CAI-1 binding, are shown. *Inset*: Zoomed-in view provides detailed visualization of these clashes.

To validate our structural observations, we introduced mutations into CqsS and examined their influence on the ability of CAI-1 to control CqsS kinase activity. We excluded TMD mutations that significantly impaired kinase activity irrespective of the presence or absence of CAI-1. Using this filter, we identified two CqsS TMD mutations that exhibit WT kinase function, CqsS W33F and CqsS F158A (Extended Data Fig. 10c,d). Consistent with the predictions from the structure, in the *in vitro* phospho-relay assay, in contrast to WT CqsS, CqsS W33F and CqsS F158A remained locked in kinase mode, transferring phosphate to LuxU under high CAI-1 levels, underscoring the essential roles that CqsS Trp33 and Phe158 play in CAI-1 sensing (Fig. 7c).

CqsS phospho-relay to LuxU requires that CqsS possesses kinase activity to catalyze the preceding autophosphorylation step. The *in vivo* bioluminescence assays show that, in *V. harveyi*, CqsS W33F and CqsS F158A do not switch from kinase to phosphatase at high cell densities, as light production is repressed under both conditions (Fig. 7d). This result demonstrates that, unlike in WT CqsS, the accumulation of endogenously produced CAI-1 does not inhibit kinase activity in CqsS W33F and CqsS F158A (Fig.7d and Extended Data Fig. 10e). Presumably, CqsS W33F and CqsS F158A fail to bind CAI-1.

### The CqsS CAI-1 sensing mechanism

To explore the CAI-1 sensing mechanism, we can compare the CqsS structures in the apo- and CAI-1-bound states. As previously mentioned, we resolved four CqsS structures lacking CAI-1 (CqsS_S1_, CqsS_S2_, CqsS-ATPγS_S1_, and CqsS-ATPγS_S2_), the TMDs of which all exhibit similar conformations (Extended Data Fig. 5f). We selected the CqsS_S1_ structure as the representative for the following comparison.

We superimposed the CqsS-CAI-1 and CqsS_S1_ structures relative to their TMDs. In each subunit, TM1-5 maintain almost unchanged conformations, with the majority of residues in the primary CAI-1 binding site undergoing only minor conformational adjustments to accommodate the CAI-1 ligand (Fig. 7e,f). When bound to CAI-1, CqsS TM6 clearly moves toward the CAI-1 ligand coordinated by the other CqsS protomer (Fig. 7e), and TM6 contributes Phe158, Phe162, and Phe166 residues to stabilize the CAI-1 molecule (Fig. 7f and Supplementary Video 2).

In the CqsS_S1_ structure, TM6 and the α1 helix of the DHp domain form a continuous elongated helix, which is crucial for maintaining the relative positions between the ICD and the TMD (Fig. 7g and Extended Data Fig. 3a). Therefore, CAI-1 binding to CqsS induces an obvious movement of TM6, potentially leading to steric clashes between TM6 Arg169 and TM1 Asn14 (Fig. 7g). Consequently, the elongated helix is disrupted at Tyr167, potentially explaining the high flexibility between the ICD and TMD observed during cryo-EM data processing (Extended Data Fig. 7d,8). Disruption of the extended helix may also interfere with the relative structural arrangements of the CA and DHp domains, thereby interfering with CqsS kinase activity.

## Discussion

CqsS is a QS receptor that fosters intra-genus cell-cell communication and is present in every sequenced vibrio species^11^. Here, our structural investigation, coupled with biochemical and genetic analyses, define the phospho-relay mechanism of CqsS and provide a structural basis for the CAI-1 sensing mechanism, thereby advancing understanding of CqsS-CAI-1-mediated QS signal transduction.

We first determined the cryo-EM structures of *V. harveyi* CqsS in apo- (CqsS_S1_ and CqsS_S2_) and ATPγS-bound (CqsS-ATPγS_S1_ and CqsS-ATPγS_S2_) states. CqsS is an asymmetric dimer, confirming an asymmetric catalytic mechanism. Our combined structural and biochemical characterization revealed *cis* autophosphorylation of the CqsS H1 residue and *trans* phospho-transfer to the CqsS D1 residue. In our ATPγS-bound structures, ATPγS is coordinated by the CA domain, with the γ-phosphate placed 3.2 Å from the H1 (His190) residue of the same subunit, mimicking the ATP-bound state required for H1 phosphorylation. To fully understand the catalytic mechanism of H1 phosphorylation, structures of proteins in additional intermediate states, such as the ADP-bound state, are necessary. Although the Rec domain of CqsS protomer A is in close proximity to the H1 residue on protomer B in the CqsS_S1_ structure, the distance between H1 and D1 (Asp613) would not allow direct phospho-transfer from H1 to D1 in the state captured in our structure (Fig. 3a and Extended Data Fig. 9b). Thus, CqsS may undergo conformational changes that bring the H1 and D1 residues in close proximity, thereby enabling the *trans* phospho-transfer. To trap the CqsS protein in this transient state, it may be necessary to develop a strategy that stabilizes phosphorylated H1 residues and inhibits subsequent phospho-transfer from H1 to D1 via mutation of D1.

We also determined the CAI-1-bound structure (CqsS-CAI-1) by co-expressing CqsS and the CAI-1 synthase CqsA. This structure provides information on how the QS autoinducer CAI-1 is coordinated by the CqsS receptor and reveals an obvious movement of TM6 induced by ligand binding (Fig. 7b,e-g). Previous genetic studies suggested that CAI-1 binds to the TMD of CqsS and binding inhibits the kinase activity of the ICD^20^. Our structural observations demonstrate that the movement of TM6 disrupts the continuous elongated helix formed by TM6 and the α1 helix of the DHp domain (Fig. 7g). One plausible hypothesis is that this disruption interferes with the structural arrangements of the CA and DHp domains, leading to the inhibition of kinase activity in the ICD. An obvious movement of the CqsS DHp domain relative to the CA domain was observed in the resolved ICD structure (CqsS-CqsA) (Extended Data Fig. 9b), whereas it is likely that CqsA binding to CqsS drives this structural rearrangement. To verify this supposition, it will be essential to determine the CqsS ICD structure when bound to CAI-1 but lacking CqsA.

Remarkably and unexpectedly, we discovered that CqsS forms a 2:4 complex with the CAI-1 synthase CqsA. The consequence of CqsS-CqsA complex formation is elimination of CqsS kinase activity, which is consistent with our structural observation that, in the CqsS-CqsA complex, the CqsS H1 residue is located far from the catalytic site in the CA domain in both protomers (Fig. 4b,c and Extended Data Fig. 9b). To the best of our knowledge, there is no previous report of a ligand synthase binding to its partner receptor and modulating the catalytic activity within the two-component histidine kinase. Thus, our discovery suggests a novel mechanism for regulation of signal transduction in the CqsA-CAI-1-CqsS pathway.

The CqsS ICD interacts with CqsA, whereas the CqsS TMD interacts with CAI-1. Both CAI-1 and CqsA binding to CqsS negatively influence CqsS kinase activity (Fig. 4b-c and 7a-d). Therefore, the CqsS receptor can be simultaneously inhibited by both its ligand and the synthase of the ligand. Each type of inhibition could offer distinct advantages in the context of signal transduction. Before CAI-1 can bind to CqsS, it needs to be produced, diffuse out of cells, accumulate in step with increasing cell density, and reach the threshold required for binding. Thus, CAI-1 inhibition of CqsS kinase should necessarily be slow and only occur at high cell density. By contrast, inhibition mediated by CqsA likely occurs more rapidly than that induced by CAI-1, owing to the direct interaction of CqsA with CqsS, which could occur immediately following the translation of each protein. *cqsS* transcription was reported to be activated by the presence of CAI-1^30^. Thus, reinforcement of CqsA mediated inhibition of CqsS could occur at high cell density. Although our structure of the CqsS-CqsA complex was determined in the presence of CAI-1, apo-CqsS also binds to CqsA (Fig. 4a). Additionally, our *in vitro* pull-down assay shows that the interaction between CqsS and CqsA is independent of CAI-1 binding (Extended Data Fig. 10f). Moreover, neither this interaction nor inhibition of CqsS kinase require CqsA to be catalytically active (Fig. 4a-c and Extended Data Fig. 7b). Thus, the two CqsS inhibitors could function independently of one another. Our current experiments do not reveal anything about the strength of kinase inhibition by CAI-1 relative to that by CqsA. Possibly, one inhibitor (i.e., CAI-1) is strong while the other (i.e., CqsA) is weak, or vice versa. Such a scenario, in which two different inhibitors of different potencies are employed alone or together, could allow fine tuning of CqsS kinase activity under different conditions.

In addition to affecting the activity of CqsS, formation of the CqsS-CqsA complex may also regulate the catalytic activity of CqsA. Such regulatory interactions have been reported between other bacterial receptors and their respective ligand synthases, and both positive and negative regulatory effects occur^31,32^. Finally, given that CqsS is located in the plasma membrane, binding to CqsA may aid in the localization of CqsA nearer to the membrane. This proximity may couple the production of the highly hydrophobic CAI-1 molecule to the membrane—its site of release from the cell. Such a mechanism could enhance CAI-1 release by minimizing loss in the cytoplasm due to adsorption to other hydrophobic cellular components.

Our work represents a major step forward in the mechanistic definition of CqsS-mediated QS signal transduction and provides the foundation for further exploration of the importance of the newly identified CqsS-CqsA complex.

## Methods

### Bacterial strains and growth conditions

*E. coli* DH5α competent cells (Alpalifebio) were used for plasmid subcloning. *E. coli* BL21 (DE3) cells (Alpalifebio) were used for the expression and purification of CqsS, CqsA, and LuxU. *E. coli* were grown in Luria-Bertani (LB) medium (1% tryptone, 1% NaCl, and 0.5% yeast extract). Unless otherwise specified, all liquid cultures were aerobically grown at 37°C. When necessary, kanamycin (50 μg/mL), ampicillin (100 μg/mL), and/or streptomycin (40 μg/mL) was added to the broth.

### Cloning of *cqsS*, *cqsA*, and *luxU*

The *V. harveyi cqsS* gene was cloned from *V. harveyi* N8T11 and inserted into the pET22b vector with C-terminal Strep and FLAG tags fused in tandem for cryo-EM sample preparation, autophosphorylation, and phospho-relay assays. To validate the *cis* phosphorylation of CqsS H1 residues and *trans* phospho-transfer from CqsS H1 to D1 residues, the *mbp* gene and 10×His tag were fused in tandem to the C-terminus of the *cqsS* gene and inserted into the pRSFDuet vector. The *cqsA* gene was also cloned with an N-terminal FLAG fusion into the pRSFDuet vector for cryo-EM sample preparation and with an N-terminal 10×His fusion into the same vector for pull-down assays. The *luxU* gene was cloned from *V. harveyi* N8T11 and inserted into the pET22b vector with an N-terminal 10×His fusion. All mutants in *V. harveyi* genes were generated via a standard two-step PCR-based strategy.

### Protein expression and purification

*E. coli* BL21 (DE3) transformed with the desired plasmid was cultured in LB medium supplemented with antibiotics at 37°C, and 1 mM isopropyl β-D-thiogalactoside (IPTG) was added when the OD_600_ reached 0.8. After further culture at 37°C for 4 h, cells were collected and resuspended in lysis buffer (25 mM Tris pH 8.0, 150 mM NaCl) supplemented with 1 mM phenylmethanesulfonylfluoride (PMSF).

#### [Purification of CqsS proteins]

Following sonication, cell debris was removed via centrifugation at 8,000 × g at 4°C for 10 min. The clarified supernatant was subjected to ultracentrifugation at 100,000 × g at 4°C for 1 h. Subsequently, the membrane pellet was resuspended in lysis buffer containing protease inhibitor cocktail (Amresco). For purification, suspensions were solubilized with 1% (w/v) lauryl maltose neopentyl glycol (LMNG, Anatrace) at 4°C for 2 h, followed by centrifugation at 20,000 × g for 1 h. Supernatants were collected and applied to Strep-Tactin® Sepharose® (IBA), followed by rinse with buffer A containing 25 mM Tris (pH 8.0), 150 mM NaCl, and 0.005% (w/v) LMNG and elution with buffer A plus 2.5 mM desthiobiotin (Macklin). Resulting eluates were loaded onto anti-FLAG M2 resin (Sigma). The resin was rinsed with buffer A, and the target protein was eluted with buffer A plus 0.2 mg/mL FLAG peptide. Eluates were concentrated and further purified by size exclusion chromatography (SEC, Superose™ 6 Increase 10/300 GL, Cytiva) in buffer A. Peak fractions were collected and concentrated to approximately 11 mg/mL for cryo-sample preparation. All CqsS variants used here were purified exclusively with Strep-Tactin® Sepharose® following the identical protocol described above.

#### [Purification of CqsS chimeras]

The CqsS-Strep and CqsS-MBP chimeras were purified via tandem affinity purification via the Strep tag on CqsS-Strep and the 10×His tag at the C-terminus of CqsS-MBP. Following sonication, the cell membrane was prepared as above. The resulting membrane proteins were extracted with 1% LMNG at 4°C for 2 h, followed by centrifugation at 20,000 × g for 1 h. Supernatants were collected and applied to Strep-Tactin® Sepharose®, followed by rinse with buffer A and elution with buffer A plus 2.5 mM desthiobiotin. Eluates were loaded onto Ni-NTA agarose (Qiagen). After rinsing with buffer A, the chimeric proteins were eluted with buffer A plus 250 mM imidazole. Eluates were concentrated and further purified by SEC in buffer A. The peak fractions were collected for phosphorylation assays.

#### [Purification of CqsS-CqsA-CAI-1 complexes]

Following sonication, the cell membrane was prepared via a protocol similar to that used for CqsS purification. The complexes were extracted with 1% LMNG at 4°C for 2 h, followed by centrifugation at 20,000 × g for 1 h. Supernatants were collected and applied to Strep-Tactin® Sepharose®. After rinsing with buffer A, complexes were eluted with buffer A plus 2.5 mM desthiobiotin. Eluates were subjected to SEC (Superose™ 6 Increase 10/300 GL, Cytiva) in buffer A. The peak fractions were collected and concentrated for cryosample preparation, which was used for structural determination of CqsS at the CqsA-bound (CqsS-CqsA) and CAI-1-bound (CqsS-CAI-1) states.

#### [Purification of CqsA proteins]

Following sonication, the cell debris was pelleted via centrifugation at 20,000 × g at 4°C for 1 h. The clarified supernatant was collected and applied to Ni-NTA agarose. After rinsing with lysis buffer plus 25 mM imidazole, the proteins were eluted with lysis buffer plus 250 mM imidazole. Eluates were subjected to ion exchange chromatography (Source™ 15Q, Cytiva) for further purification.

#### [Purification of LuxU protein]

Following sonication, cell debris was pelleted by centrifugation at 20,000 × g at 4°C for 1 h. The supernatant was collected and applied to Ni-NTA agarose resin. After rinsing with buffer B (50 mM Tris pH 8.0, 200 mM NaCl, 5 mM *β*-mercaptoethanol, and 5 mM MgCl_2_) plus 20 mM imidazole, LuxU was eluted with buffer B plus 250 mM imidazole. The eluate was subjected to ion exchange chromatography and SEC (Superdex™ 200 Increase 10/30 GL, Cytiva) for further purification.

### Cryo-EM sample preparation

Cryo-EM grids were prepared via Thermo Fisher Vitrobot Mark IV. Quantifoil R1.2/1.3 Cu grids were glow-discharged with air in a PDC-32G-2 plasma cleaner (Harrick) with mid-force for 85 s. Aliquots of 3.5 μL of purified CqsS or CqsS-CqsA complex, concentrated to approximately 11 mg/mL, were applied to the glow-discharged grids. After blotting with filter paper for 3.5 s (100% humidity and 8°C), the grids were plunged into liquid ethane cooled with liquid nitrogen. To capture the ATPγS-bound state, CqsS was combined with 5 mM ATPγS and 10 mM MgCl_2_ on ice for 1 h before grid preparation.

Grids were loaded into a Titan Krios (FEI) electron microscope operating at 300 kV equipped with a BioQuantum energy filter and a K3 direct electron detector (Gatan). Images were automatically collected with EPU in super-resolution mode. Defocus values varied from -1.5 to -1.7 μm. Image stacks were acquired with an exposure time of 3.8 s and fractionated into 32 frames with a total dose of 50 e^-^ Å^−2^. The stacks were motion corrected with MotionCor2^33^ and binned two-fold, resulting in a pixel size of 0.82 Å/pixel; meanwhile, dose weighting was performed^34^. The defocus values were estimated via patch CTF estimation.

### Cryo-EM data processing

In total, 1,630, 1,434, and 2,731 micrographs were collected for CqsS, CqsS-ATPγS, and CqsS-CqsA-CAI-1, respectively. After the particles were picked, 1,492,169, 1,523,166, and 1,679,523 particles were extracted for further data processing, respectively. According to the micrographs of the CqsS-CqsA-CAI-1 complex, two EM structures corresponding to the ICD and TMD were determined, which were designated CqsS-CqsA (ICD structure) and CqsS-CAI-1 (TMD structure) in the main text.

#### [CqsS]

The particles were applied to 2D classification, resulting in 188,821 good particles. After ab-initio reconstruction with 4 classes, one good class was outstanding. This good class served as reference for heterogeneous refinement, yielding good particles for another run of ab-initio reconstruction. The resulting good class, containing 271,340 particles, was selected as seeds for seed-facilitated classification to select additional good particles from the original particles, resulting in 713,012 good particles. These good particles were further subjected to heterogeneous refinement, resulting in 356,995 good particles and a 3.08 Å 3D reconstruction. After 3D classification with a mask on the ICD, two distinct classes were selected, Class A with 124,184 particles and Class B with 137,177 particles, yielding 3D reconstitutions of 3.24 Å and 3.28 Å, respectively. These two classes were designated CqsS_S1_ and CqsS_S2_, respectively.

#### [CqsS-ATPγS]

After 2D classification, a total of 381,945 good particles were selected and subjected to ab-initio reconstruction, resulting in a good class with 227,283 particles. These particles were selected as seeds for seed-facilitated classification, yielding 580,605 good particles. After a further 3D classification with a mask on the ICD, two distinct classes were selected, Class A with 150,052 particles and Class B with 140,065 particles, yielding 3D reconstitutions with 3.48 Å and 3.46 Å, respectively. These two reconstructions were designated CqsS-ATPγS_S1_ and CqsS-ATPγS_S2_, respectively.

#### [CqsS-CqsA-CAI-1]

After 2D classification, 2D averages could be separated into two major classes: averages with stronger densities in the ICD and those with stronger densities in the TMD. These two types of 2D averages were chosen as template 1 and template 2 to select additional particles from the original micrographs with estimated particle diameters of 170 Å and 120 Å, resulting in 1,855,902, and 3,415,200 particles, respectively. These particles were applied to the structural determination of the ICD and TMD of this complex.

For the ICD, 1,855,902 particles were subjected to several rounds of 2D classification, resulting in a total of 318,201 good particles. These particles were used in ab-initio reconstruction, resulting in good reconstruction with 200,271 particles. These particles were selected as seeds for seed-facilitated classification, yielding 470,547 good particles. After further 3D classification, the ICD of CqsS-CqsA was determined at 2.87 Å resolution with 130,780 particles. This structure is designated CqsS-CqsA in the main text.

For the TMD, 3,415,200 particles were subjected to 2D classification, heterogeneous refinement, and ab initio reconstruction, resulting in a good reconstruction with 239,375 particles. These particles were selected as seeds for seed-facilitated classification, yielding 737,497 good particles. After several rounds of heterogeneous refinement and 3D classification, 131,035 good particles were selected which yielded a 3D reconstruction at 2.87 Å resolution after non-uniform refinement with a mask on the TMD. This structure is referred to as CqsS-CAI-1 in the main text.

### Model building and refinement

The structure of CqsS was built primarily based on the CqsS_S1_ map at 3.24 Å resolution. AlphaFold2-predicted models^35^ facilitated model building for regions with moderate resolution. The resulting structure served as an initial reference for model building of CqsS_S2_, CqsS-ATPγS_S1_, CqsS-ATPγS_S2_, CqsS-CqsA, and CqsS-CAI-1. Manual adjustment was made in Coot^36^, and all structure refinement was carried out by PHENIX^37^ in real space with secondary structure and geometry restraints. All the structure figures were prepared with PyMol^38^, and all the EM maps were prepared with Chimera^39^.

### *In vitro* CqsS phosphorylation assay

Phosphorylation assays were conducted in phosphorylation buffer (50 mM Tris pH 8.0, 100 mM KCl, 5 mM MgCl_2_, and 10% (v/v) glycerol) with 2 μM CqsS, CqsS mutants, or CqsS chimeras. For the assays involving LuxU, LuxU was added at 10 μM. For the CqsA inhibition experiments, 2 μM CqsA was included. For the CAI-1 inhibition experiments, CAI-1 was produced by overexpressing CqsA in *E. coli* BL21 cultured in LB medium as previously reported ^10^, and cell-free fluid was added to the assay system 30 min before the reactions were initiated. All experiments were initiated by adding 100 μM cold ATP and 0.5 μCi [γ-^32^P]-ATP (from a stock of 3,000 Ci/mmol, Revvity). To verify *cis* phosphorylation on CqsS H1, only 0.5 μCi [γ-^32^P]-ATP was used to initiate the reaction, and no cold ATP was added. After incubation at room temperature for 10 min, the reactions were terminated with SDS loading buffer. Following separation via SDS-PAGE, gels were exposed to a phosphoscreen overnight and subsequently analyzed via Amersham™ Typhoon™ IP and ImageJ.

### *In vivo* bioluminescence assay

WT *V. harveyi* N8T11 *cqsA* and *cqsS* sequences were subcloned into pMMB, together with the corresponding *cqsAS* upstream and downstream regulatory regions from *V. harveyi* BB120. This construct was used as the template to generate the desired *cqsS* mutations via site-directed mutagenesis. To test the effects of mutations on CqsS function, the different versions of *cqsAS* along with the upstream and downstream *V. harveyi* BB120 regulatory DNA were amplified via PCR and inserted into the pLAFR2 cosmid via the BamHI and XbaI restriction sites. The resulting constructs were introduced into *V. harveyi* WN1397 (BB120 *ΔluxN ΔluxPQ ΔcqsAS*) by conjugation. Exconjugate strains were grown overnight in LM at 30°C with shaking, followed by back-dilution to OD_600_ = 0.005. Aliquots (150 µL) of the cultures were transferred into the wells of 96-well plates (Corning). The plates were incubated at 30°C (Biotek Biospa). The plates were shaken every 15 min. Light production and OD_600_ were measured (Biotek Neo). Light output is reported as relative light units (RLUs), which is the bioluminescence/OD_600_.

### CqsS-CqsA pull-down assay

0.2 mg of His-tagged CqsA and 0.1 mg of Strep-tagged CqsS were incubated in 1 mL of buffer A on ice for 1 h. Subsequently, the mixtures were loaded onto 300 μL of Strep-Tactin® Sepharose®. After washing with buffer A, the resin was eluted with buffer A supplemented with 2.5 mM desthiobiotin. Eluates were subjected to SDS‒PAGE analysis. To evaluate the effect of CAI-1 on the interaction, cell-free culture fluids from *E. coli* expressing CqsA or the inactive CqsA K240A variant were included during the incubation period.

### Statistics and reproducibility

No statistical methods were used to predetermine sample sizes. The experiments were not randomized, and the investigators were not blinded during the experiments nor outcome assessments. Each experiment was conducted independently at least three times with similar results.

### Data availability

The plasmids generated in this study will be made available upon request. The atomic coordinates of CqsS_S1_, CqsS_S2_, CqsS-ATPγS_S1_, CqsS-ATPγS_S2_, CqsS-CqsA, and CqsS-CAI-1 have been deposited in the PDB (http://www.rcsb.org) under the accession codes 9WXL, 9WXQ, 9WY6, 9WY7, 9WY4, and 9WXU, respectively. The corresponding electron microscopy maps have been deposited in the Electron Microscopy Data Bank (https://www.ebi.ac.uk/pdbe/emdb/) under the accession codes EMD-66349, EMD-66354, EMD-66361, EMD-66365, EMD-66360, and EMD-66357.

### Code availability

No custom code or mathematical algorithm was used in this study.

## Acknowledgments

We thank the Cryo-EM Center of the University of Science and Technology of China (USTC) for EM facility support. We are grateful to Dr. Yong-Xiang Gao and all the other staff members at the Cryo-EM Center for their technical support in cryo-EM data collection. We thank Dr. Zhen Tao from Zhejiang Ocean University for sharing the *V. harveyi* N8T11 strain. H.Q. was supported by the National Key R&D Program of China (2024YFA1803104 & 2024YFA1307900), the National Natural Science Foundation of China (32271241), and start-up funding from the University of Science and Technology of China (KY9100000034 and KJ2070000082). B.L.B was supported by the Howard Hughes Medical Institute, the National Institutes of Health grant R37GM065859 and the National Science Foundation grant MCB-2508324. G.A.B. was supported by NIH Grant F32GM149034.

## Author contributions

H.Q. and M.L. conceived the project and designed the experiments. M.L., Y.Y., and Y.Z. performed cloning and protein purification. Y.Y. prepared the cryo-EM samples and collected the data. H.Q. determined the structures. Y.Y., M.L., and Y.Z. performed the *in vitro* phosphorylation assays. Y.Y. and M.L. performed the *in vitro* pull-down assays. J.S.V. and G.A.B. cloned the genes required for and performed the *in vivo* bioluminescence assays. H.Q., M.L., Y.Y., B.L.B., and J.S.V. analyzed the data. H.Q., B.L.B., M.L., and Y.Y. wrote the initial draft, and all authors contributed to the revision of the manuscript.

