## Extended data Fig. 1-10; Supplementary Table 1; Supplementary Video 1-2 for "Molecular basis of quorum-sensing signal transduction by CqsS and its inhibition by CqsA the autoinducer synthase": Extended Data.pdf

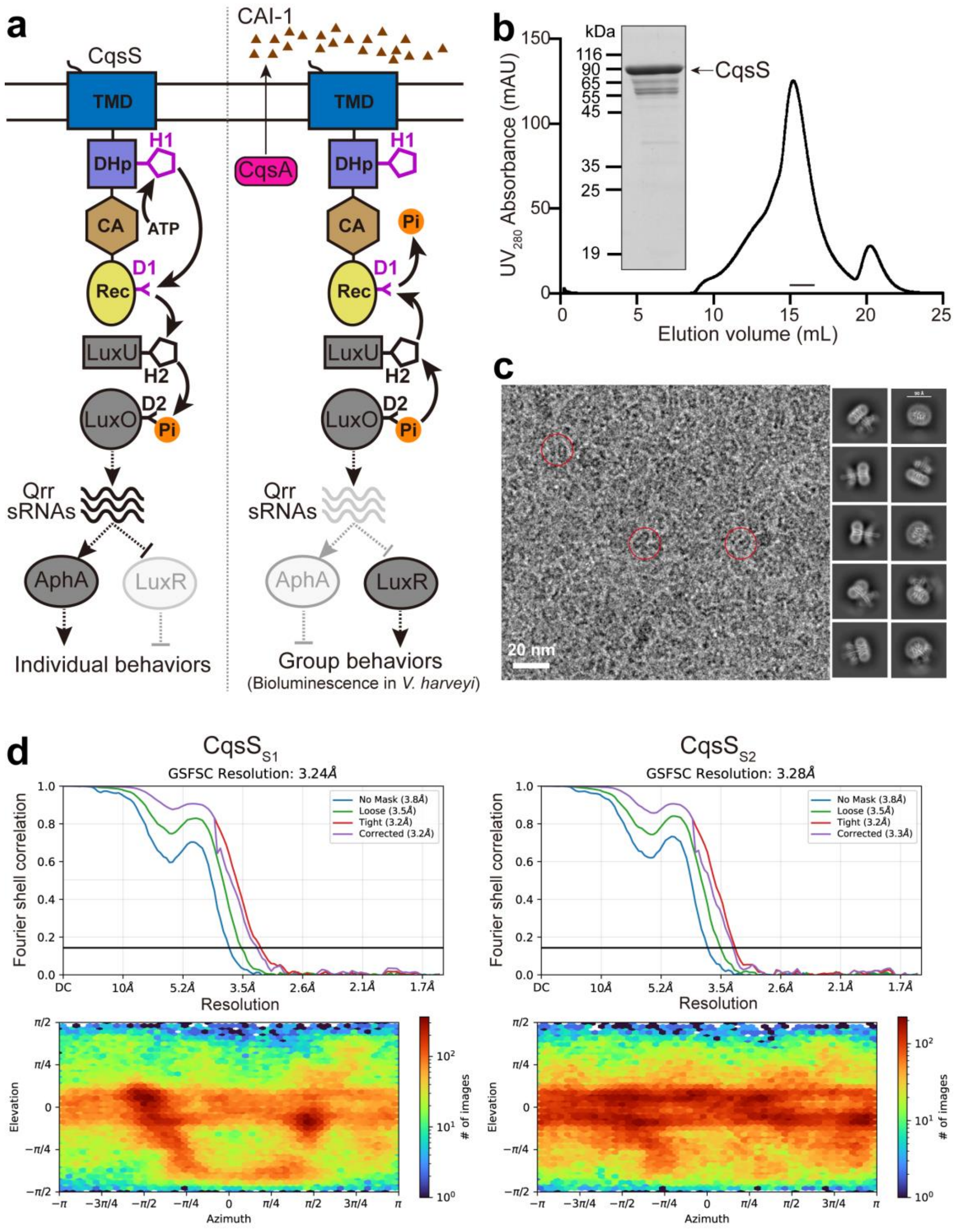


**Extended Data Fig. 1 | Cryo-EM analysis of *V. harveyi* CqsS. a**, A schematic diagram illustrating CqsS-mediated QS signal transduction under low (left) and high (right) cell density conditions in vibrios. The brown triangles represent the CAI-1 autoinducer. **b**, Representative size exclusion chromatography (SEC) profile of CqsS solubilized in

0.005% LMNG. Peak fractions were concentrated for cryo-EM sample preparation. *Inset*: SDS-PAGE analysis of the concentrated CqsS protein used for cryo-EM sample preparation. The experiments were independently repeated more than three times with similar results. **c**, Representative micrographs (left) and 2D class averages (right) of cryo-samples of CqsS. The representative particles are indicated by red circles. Scale bar, 20 nm. **d**, Golden standard Fourier shell correlation (GSFSC) curves (top) and angular distribution maps (bottom) for cryo-EM reconstructions of $CqsS_{S1}$ (left) and $CqsS_{S2}$ (right).

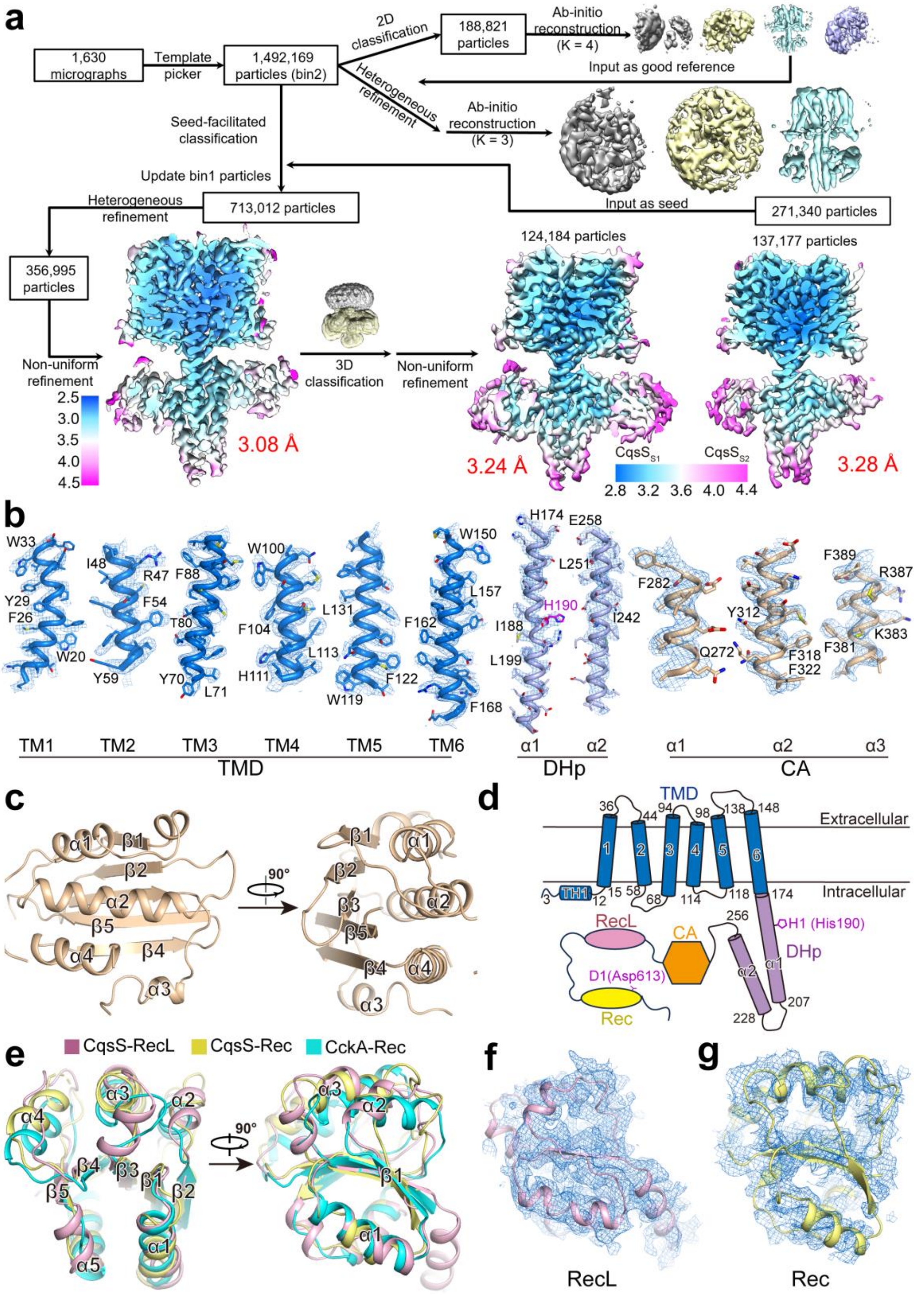


**Extended Data Fig. 2 | Structural determination and model building of CqsS. a**, Flowchart for the structural determination of CqsS in two states, $CqsS_{S1}$ and $CqsS_{S2}$. The unit of resolution is Å. Details are provided in the Methods section. **b**, EM maps of representative structural elements. The densities, contoured at 5-6σ, were prepared in PyMOL based on the $CqsS_{S1}$ map. **c**, The structure of the CqsS CA domain. This domain consists of 4 helices and 5 sheets. **d**, Membrane topology of CqsS based on the cryo-EM structures. **e**, Structural overlay of the RecL and Rec domains of CqsS relative to the Rec domain from the representative hybrid histidine kinase CckA (PDB code: 6TNE). **f**, **g**, EM maps of the RecL (**f**) and Rec (**g**) domains of CqsS. All figures were generated based on the CqsS-A protomer in the $CqsS_{S1}$ structure.

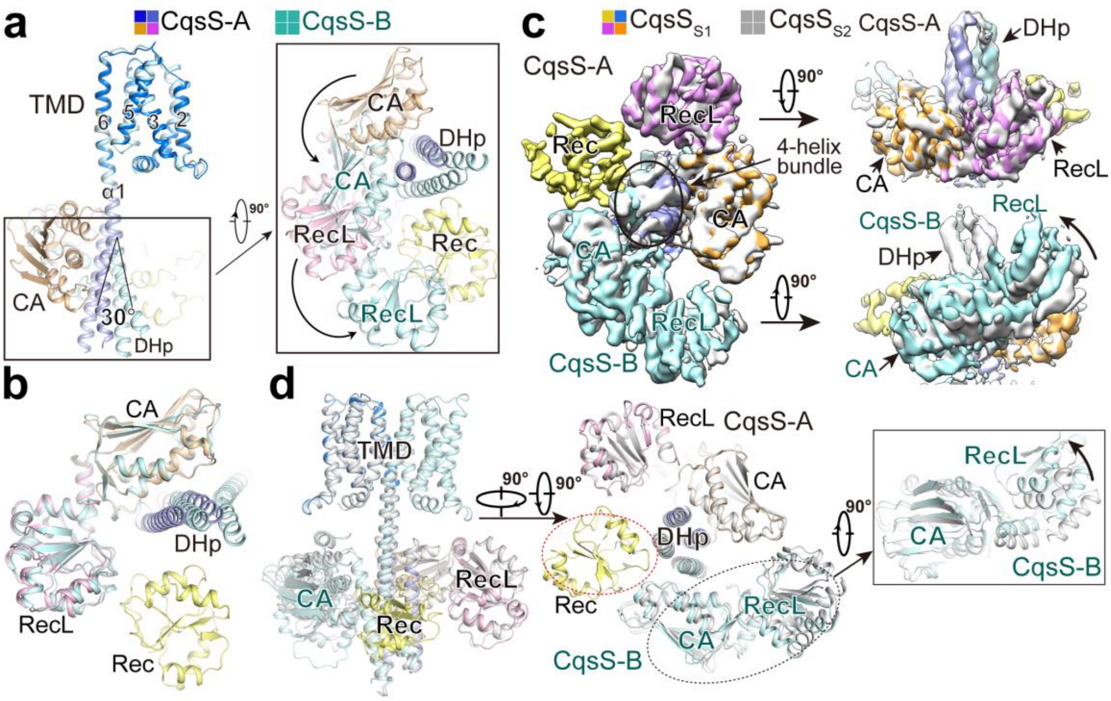


**Extended Data Fig. 3 | Structural comparison of CqsS$_{S1}$ and CqsS$_{S2}$. a**, Structural superimposition of two protomers (CqsS-A and CqsS-B) in the CqsS$_{S1}$ structure relative to their TMDs. *Inset*: Zoomed-in view highlighting structural differences in the cytosolic domains between the two protomers. **b**, A cytosolic view showing structural comparison between the CqsS-A and CqsS-B protomers relative to their CA domains. **c**, Superimposition of the EM maps of CqsS$_{S1}$ and CqsS$_{S2}$ to reveal differences in their cytosolic domains (left), and two separate views (right) show the differences in each protomer: CqsS-A (top) and CqsS-B (bottom). **d**, One membrane view and one extracellular view to compare the CqsS$_{S1}$ and CqsS$_{S2}$ structures. The Rec domain of CqsS-A is not visible in the CqsS$_{S2}$ structure, as indicated by the red ellipse, and obvious structural differences are highlighted in the CA and RecL domains of CqsS-B by the black ellipse. *Inset*: Zoomed-in view showing the conformational changes that exist in the CA and RecL domains of CqsS-B between these two structures.

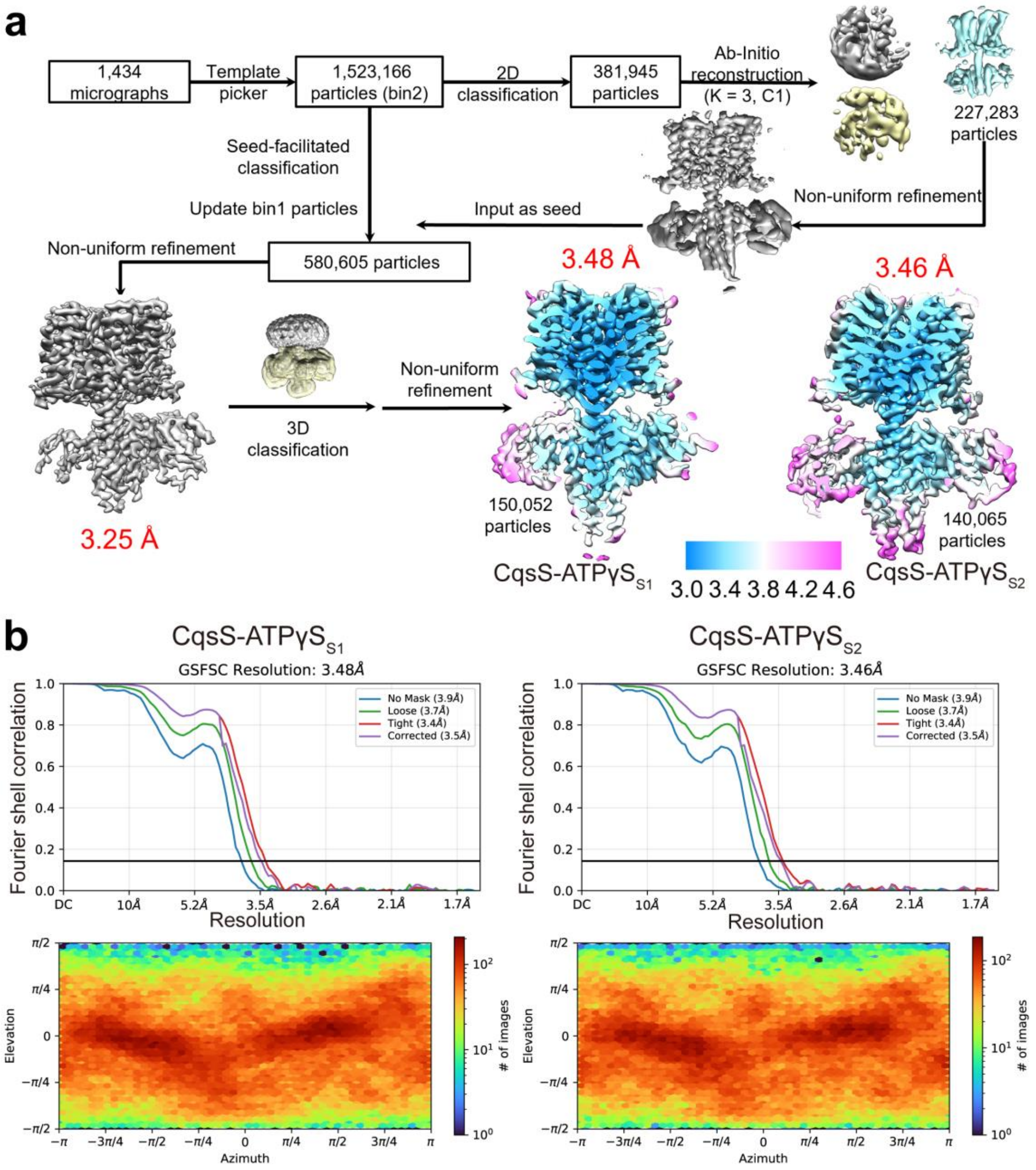


**Extended Data Fig. 4 | Structure determination of CqsS in the ATP γ S-bound state.**

**a**, Flowchart for the structural determination of CqsS-ATPγS complexes in two states, CqsS-ATPγS$_{S1}$ and CqsS-ATPγS$_{S2}$. The unit used for resolution is Å. **b**, GSFSC curves (top) and angular distribution maps (bottom) for CqsS-ATPγS$_{S1}$ (left) and CqsS-ATPγS$_{S2}$ (right).

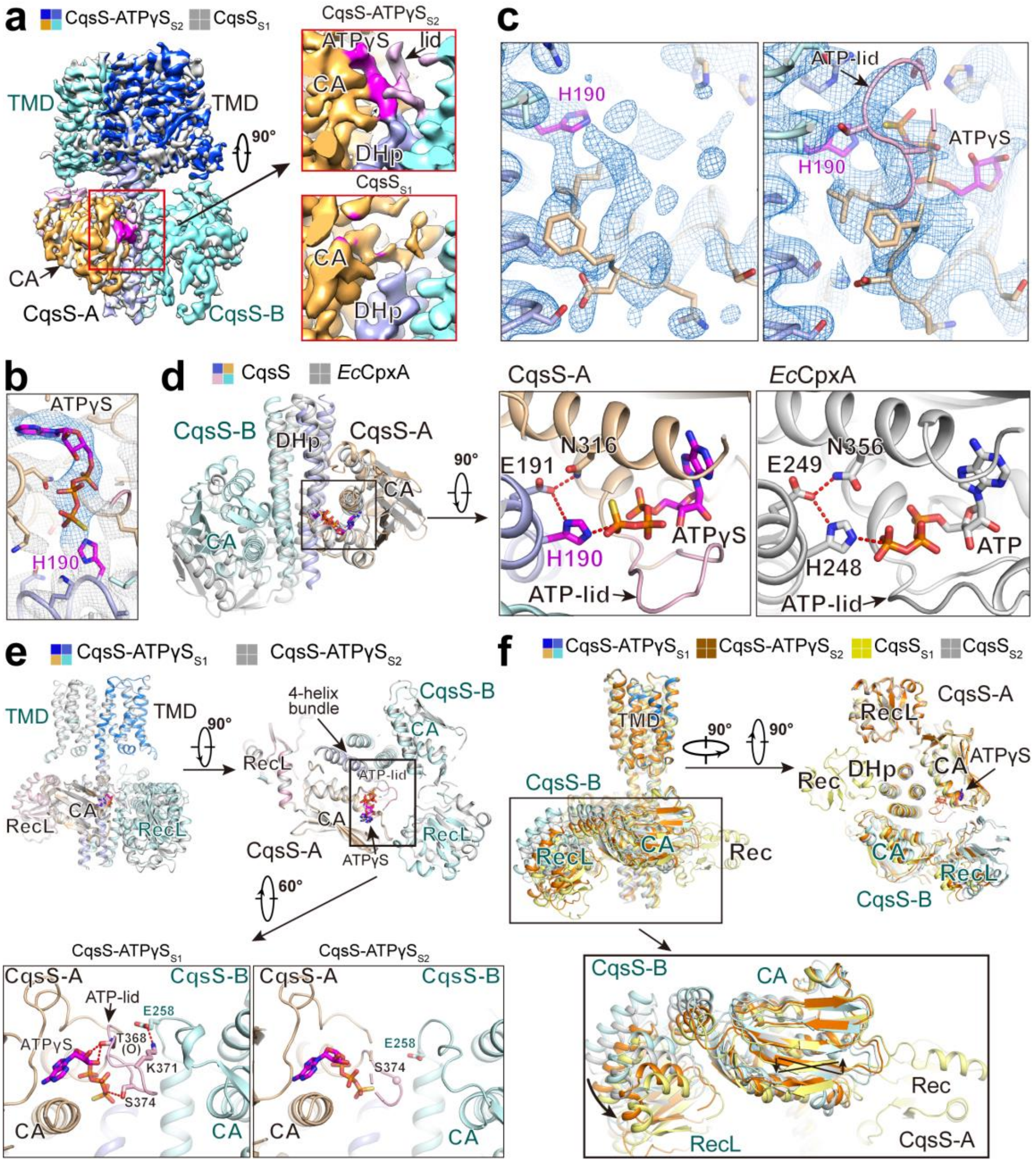


**Extended Data Fig. 5 | Structural comparison of CqsS in the apo- and ATPγS-bound states. a**, Overlay of maps between CqsS$_{S1}$ and CqsS-ATPγS$_{S2}$. *Insets*: Zoomed in views showing the ATP binding site in the CA domain of the CqsS-A protomer in CqsS-ATPγS$_{S2}$ (top) and CqsS$_{S1}$ (bottom). **b**, Structural modeling of the ATPγS molecule into the densities in the CqsS-ATPγS$_{S2}$ structure. **c**, Comparative views of EM maps at the ATP-lid region between CqsS$_{S1}$ (left) and CqsS-ATPγS$_{S2}$ (right). The

EM maps for this region of CqsS-ATPγS$_{S1}$ are shown in Fig. 2c. All the EM maps are contoured at 6σ. **d**, Structural comparison between the histidine kinase domains (DHp and CA domains) of CqsS and the classic histidine kinase domain of *E. coli* CpxA (PDB code, 5LFK). *Insets*: Zoomed-in views detailing the interactions between the CA and DHp domains in these two proteins. **e**, Structural comparison between CqsS-ATPγS$_{S1}$ and CqsS-ATPγS$_{S2}$. Top: A membrane view (left) and an extracellular view (right) are presented to show the overlay of the two structures. Bottom: Zoomed in views showing the interaction between two CA domains in the two structures. A salt bridge between CqsS-A Lys371 and CqsS-B Glu258 is observed in the CqsS-ATPγS$_{S1}$ structure (left) but not in the CqsS-ATPγS$_{S2}$ structure (right), potentially explaining the partial visualization of the ATP-lid loop in CqsS-ATPγS$_{S2}$. **f**, Structural comparison between CqsS$_{S1}$, CqsS$_{S2}$, CqsS-ATPγS$_{S1}$, and CqsS-ATPγS$_{S2}$. Top: One membrane (left) and one cytosolic (right) view showing structural overlays of these four structures. The TMDs are nearly identical, while some obvious structural differences are observed in the cytosolic domains, primarily in the CqsS-B protomer. Bottom: Zoomed in view showing the conformational changes in the CA and RecL domains of CqsS-B among these structures.

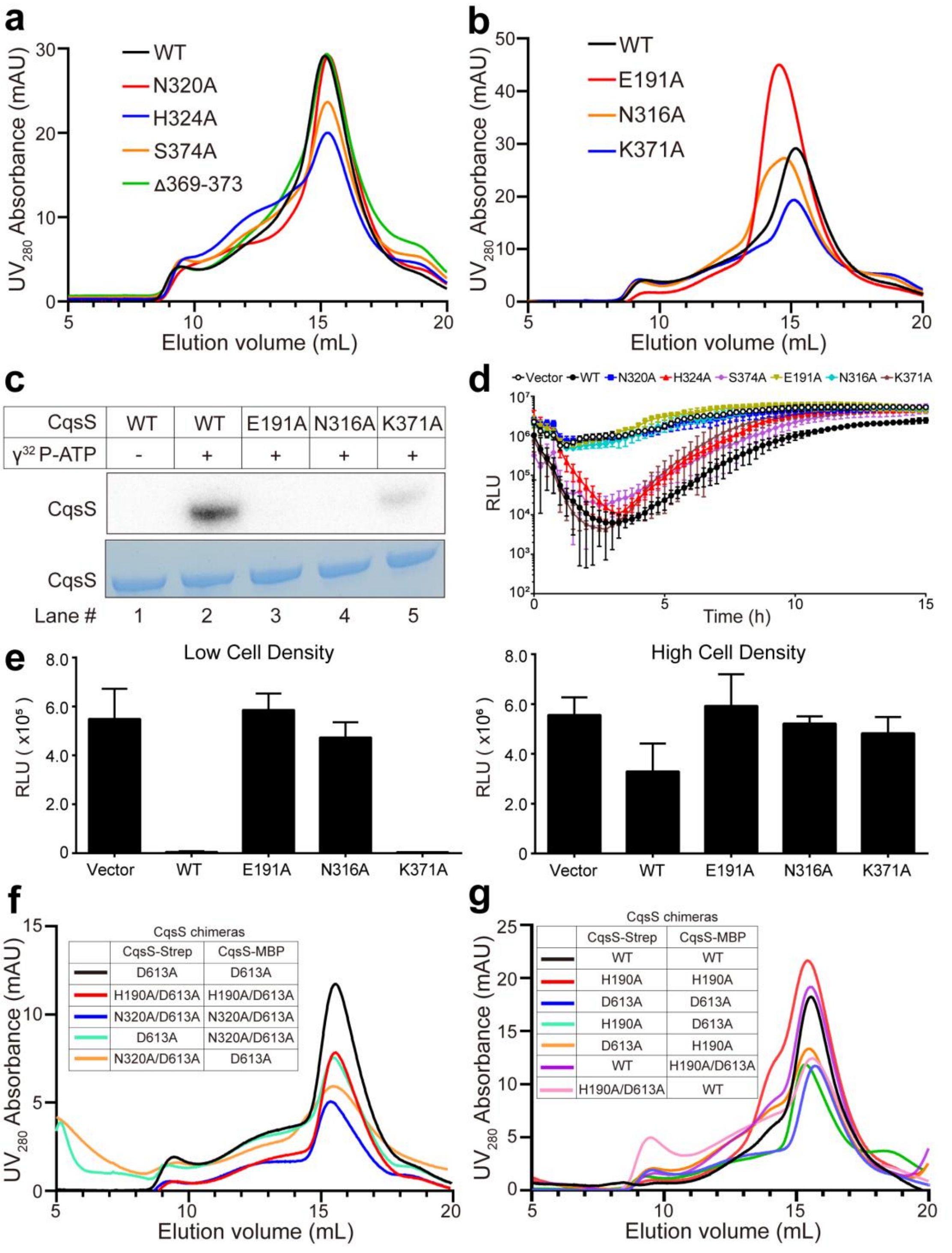


**Extended Data Fig. 6 | SEC profiles of WT CqsS, CqsS variants, and CqsS chimeras used in phosphorylation assays. a**, SEC profiles of WT CqsS and CqsS variants investigated for ATPγS coordination. **b**, SEC profiles of WT CqsS and CqsS variants investigated for interactions between the CA and DHp domains. The CqsS variants in panels **a** and **b** have elution volumes comparable to that of the WT CqsS, supporting their use in the *in vitro* autophosphorylation assays shown in Fig. 2f and Extended Data Fig. 6c. **c**, Validation of the essential residues involved in the CqsS CA-DHp interaction as judged by the CqsS autophosphorylation assay. Purified WT and variant CqsS proteins were incubated with radioactive ATP ($\gamma^{32}$P-ATP) in phosphorylation buffer, followed by autoradiography (top, gray) and Coomassie blue staining (bottom, blue). **d**, The bioluminescence output of Δ*cqsS V. harveyi* was assessed continuously over time following the introduction of the designated *cqsS* mutants. The minimum and maximum RLU values were selected to generate the bar graphs for low and high cell densities presented in Fig. 2g and Extended Data Fig. 6e, respectively. See the Methods section for all assay conditions. **e**, Validation of the essentiality of the designated residues involved in intra-domain interactions via assessment of *in vivo* kinase activity in Δ*cqsS V. harveyi* carrying different CqsS variants. The graphs present the minimum and maximum light output in the assay, which occur at low cell density (left) and high cell density (right), respectively. In panels **d** and **e**, RLU refers to the relative light units, which are the bioluminescence per $OD_{600}$. Error bars denote standard deviations of the means, *n*=3. **f**, **g**, SEC profiles of CqsS-Strep and CqsS-MBP chimeras used to verify *cis* phosphorylation on the CqsS H1 residue (**f**) and *trans* phospho-transfer from the H1 to D1 residue (**g**). The mutations present in each subunit of the CqsS chimeras are detailed in the accompanying tables. See Methods for details.

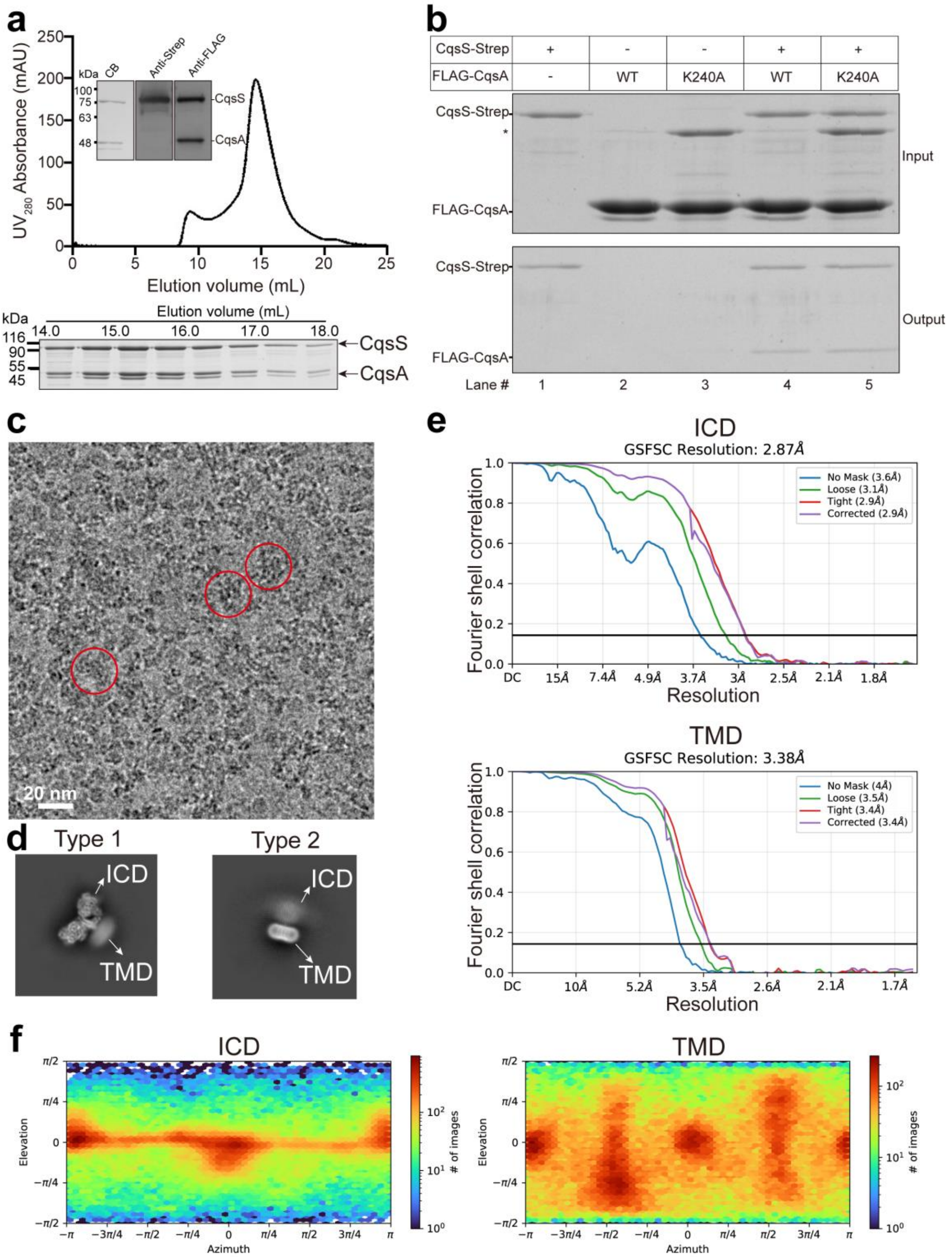


**Extended Data Fig. 7 | Cryo-EM analysis of the CqsS-CqsA-CAI-1 complex. a**, Representative SEC profile of the CqsS-CqsA-CAI-1 complex. *Insets*: SDS-PAGE analysis of the peak fractions, followed by Coomassie blue analysis (left) and immunoblotting analyses using anti-Strep (middle) and anti-FLAG (right) antibodies. The peak fractions were concentrated for cryo-EM sample preparation (bottom). The experiments were independently repeated more than three times with similar results. **b**, The catalytically-inactive CqsA K240A mutant protein interacts with CqsS with comparable avidity as WT CqsA. WT CqsA and CqsA K240A were incubated with CqsS followed by affinity purification using a Strep tag on CqsS. See Methods for details. **c**, Representative micrograph of a cryo-EM sample of the CqsS-CqsA-CAI-1 complex. The representative particles are indicated by red circles. Scale bar, 20 nm. **d**, Typical images of two distinct types of 2D averages. Left: 2D average with a clear intracellular domain (ICD) but a blurred TMD; right: 2D average with a clear TMD but a blurred ICD. **e**, GSFSC curves for the final reconstructions of the ICD (top, CqsS-CqsA) and TMD (bottom, CqsS-CAI-1) for the CqsS-CqsA complex. **f**, Angular distribution maps for the final reconstructions of the ICD (left, CqsS-CqsA) and TMD (right, CqsS-CAI-1) for the CqsS-CqsA complex. The ICD and TMD structures are referred to as the CqsS-CqsA and CqsS-CAI-1 structures, respectively, in the main text.

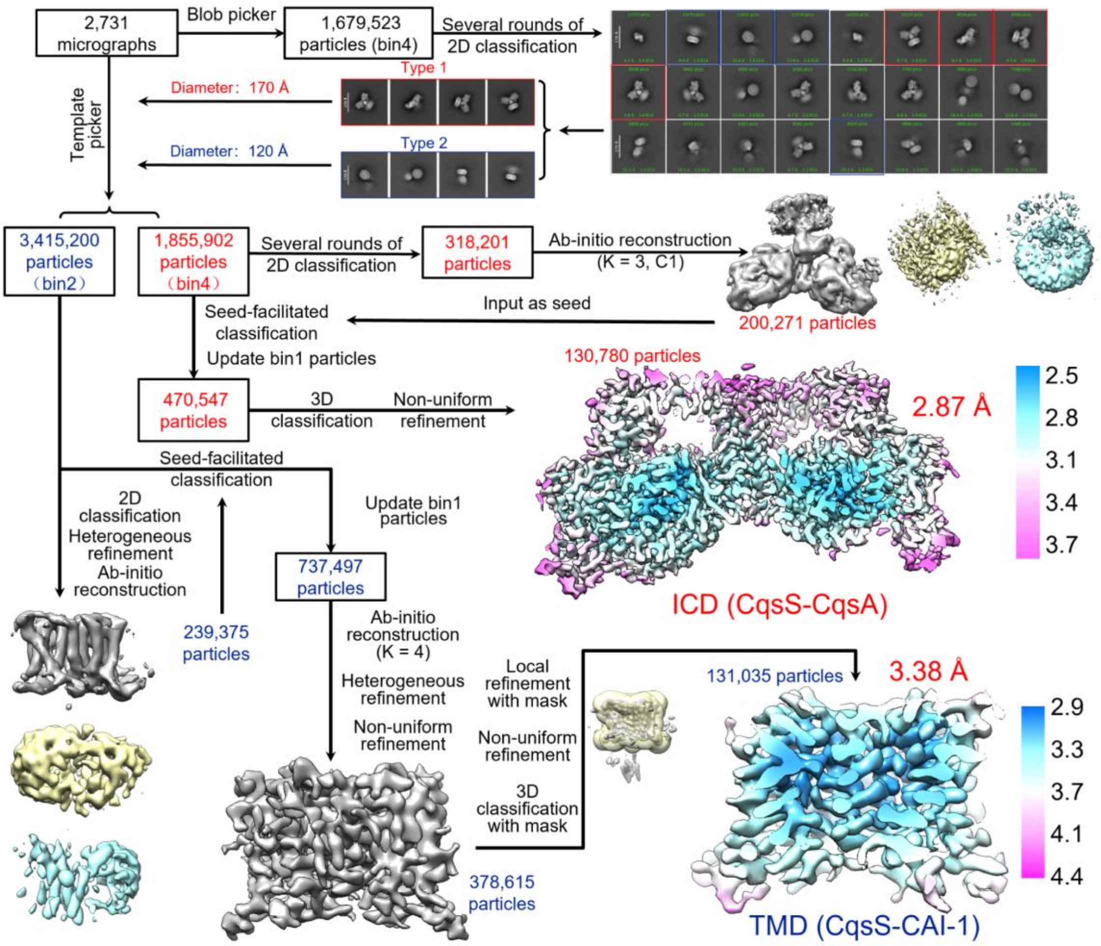


**Extended Data Fig. 8 | Structural determination of the CqsS-CqsA-CAI-1 complex.**

Flowchart for the structural determination of the ICD and TMD of the CqsS-CqsA-CAI-1 complex. The unit used for resolution is Å. Red and blue text indicate the structural determination of the ICD and TMD, which are referred to as the CqsS-CqsA and CqsS-CAI-1 structures in the main text, respectively. See Methods for details.

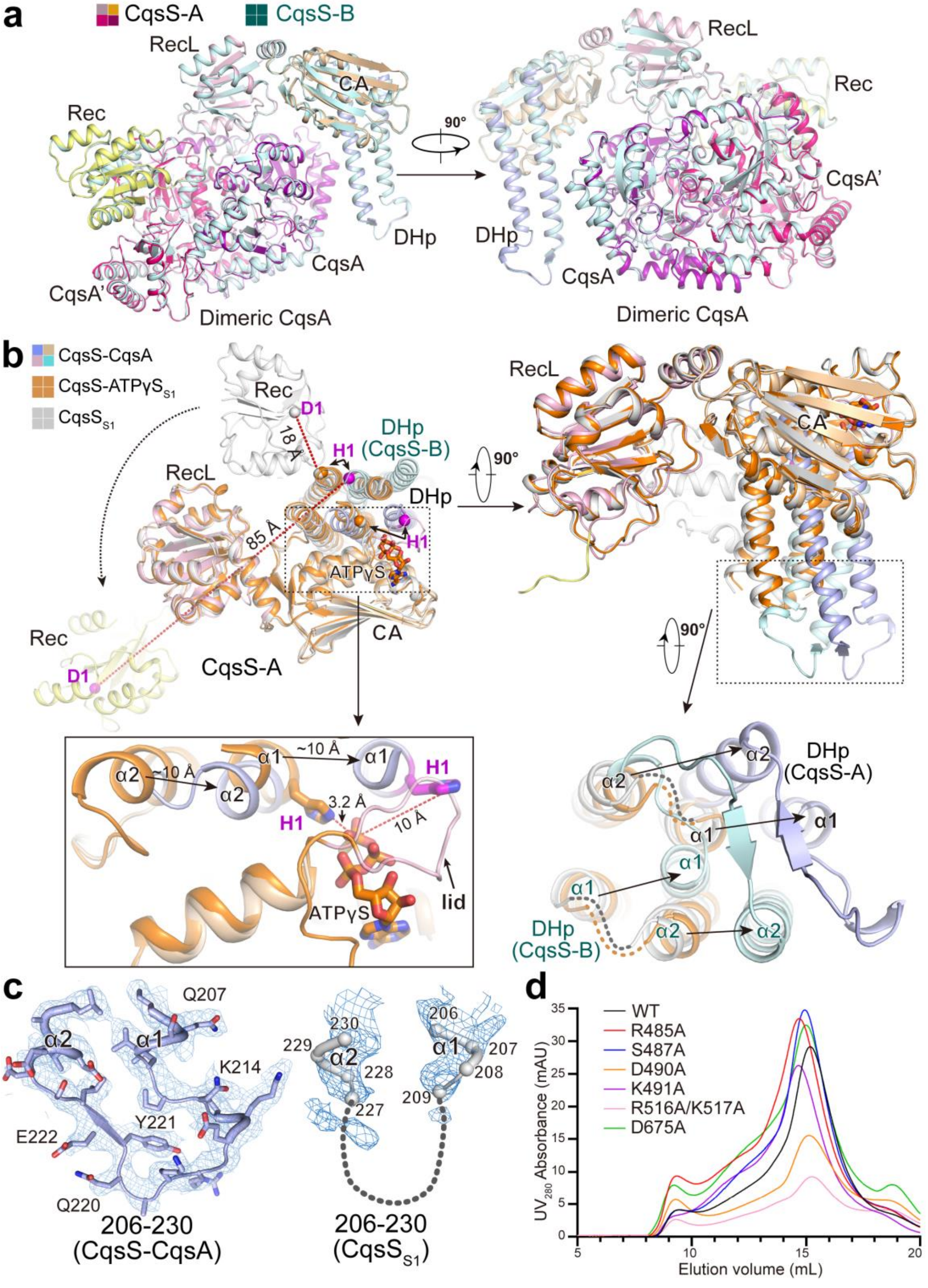


**Extended Data Fig. 9 | Structural comparison among CqsS-CqsA, CqsS-ATPγS$_{S1}$, and CqsS$_{S1}$. a**, Two views showing superimpositions of two 1:2 CqsS-CqsA subcomplexes, suggesting high similarity. **b**, Structural overlay of the CqsS-A protomer in the CqsS-CqsA, CqsS-ATPγ$S_{S1}$, and CqsS$_{S1}$ structures. The DHp domain of CqsS-B is included to illustrate the distance between the H1 residue of CqsS-B and the D1 residue of CqsS-A, where a *trans* phospho-transfer may occur. Top: Two views displaying the structural superimposition of the cytosolic domains of CqsS-A in relation to the CA and RecL domains for these structures. Bottom: Zoomed-in views showing the ATP binding pocket within the CA domain (left) and highlighting the structural differences between the DHp domains in these structures (right). **c**, EM maps for the loops (residues 206-230) connecting the α1 and α2 helices of the DHp domains in the CqsS-CqsA complex (left) and CqsS$_{S1}$ (right) structures. In the CqsS-CqsA complex, this loop is well-resolved, whereas it is poorly resolved in the CqsS$_{S1}$ structure. The EM maps are contoured at $5\sigma$. **d**, SEC profiles of CqsS mutants with substitutions in residues involved in interaction with CqsA. The CqsS variants have elution volumes similar to that of the WT CqsS.

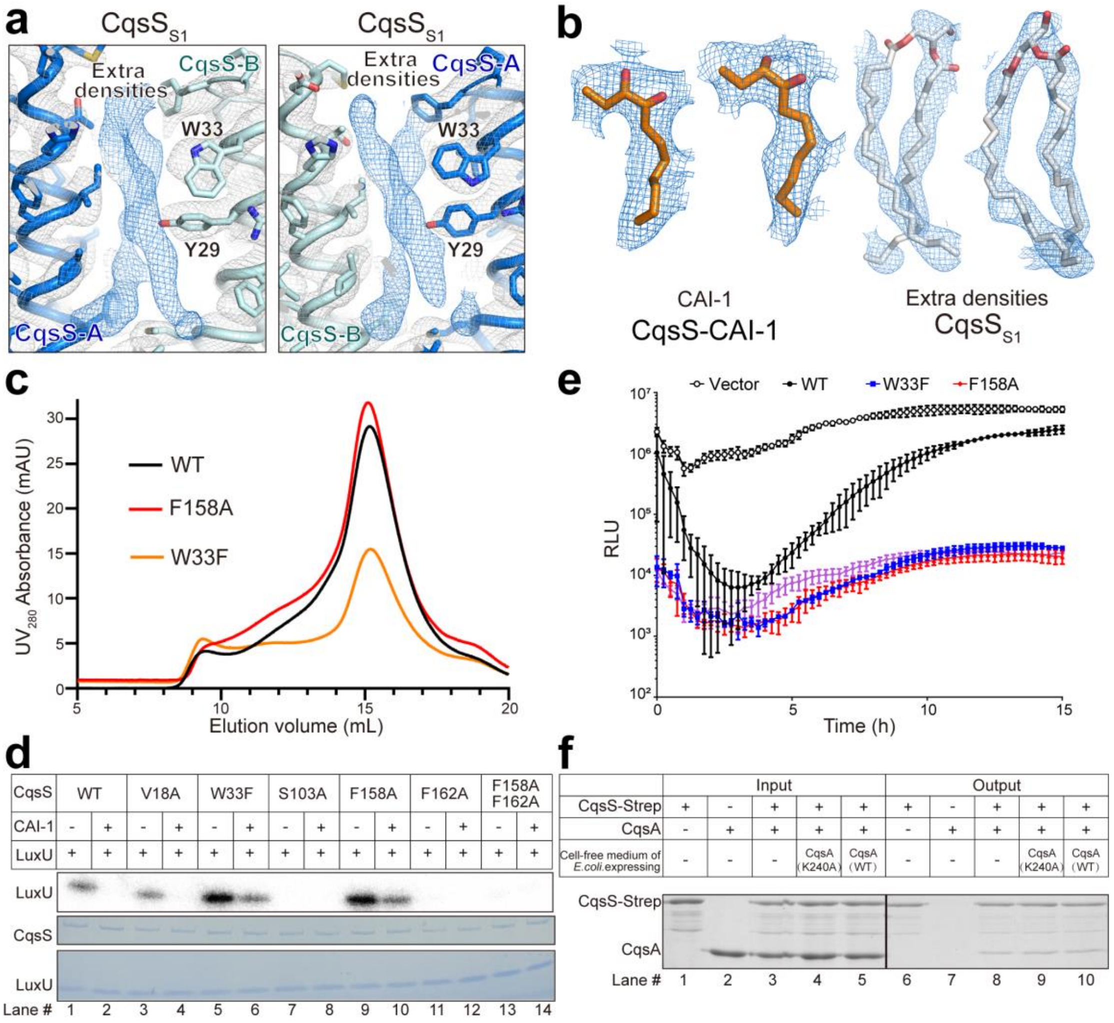


**Extended Data Fig. 10 | Additional densities at the CAI-1 binding sites in the CqsS$_{S1}$ structure and validation of essential residues involved in CAI-1 coordination and signal transduction. a**, The extra densities at the CAI-1 binding sites in the CqsS$_{S1}$ structure are contoured at 6σ. For comparison, please refer to Fig. 6c for the CAI-1 densities in the CqsS-CAI-1 complex. **b**, EM maps for the CAI-1 molecules in CqsS-CAI-1 complex (left) and the extra densities at the corresponding sites in CqsS$_{S1}$ (right) shown with diacylglycerol. The EM maps are contoured at 6σ. **c**,

SEC profiles of CqsS variants used to validate the residues that participate in CAI-1 coordination. The CqsS variants have elution volumes similar to that of the WT CqsS. **d**, *In vitro* assessment of the kinase activities of CqsS variants related to CAI-1 coordination. Purified CqsS proteins were incubated with radioactive ATP ($\gamma^{32}$P-ATP) and purified LuxU protein in the presence or absence of a cell-free culture medium containing CAI-1 (5% of the original cell-free culture fluid), followed by autoradiography (top, gray) and Coomassie blue staining (bottom, blue). **e**, Bioluminescence output of Δ*cqsS V. harveyi* was assessed continuously over time following the introduction of *cqsS* W33F and *cqsS* F158A. RLU refers to relative light units, which are the bioluminescence per $OD_{600}$. Error bars denote standard deviations of the means, *n*=3. The minimum and maximum RLU values were selected to generate the bar graphs for low and high cell densities, respectively, as shown in Fig. 7d. **f**, Assessment of the influence of CAI-1 on the direct interaction between CqsS and CqsA via *in vitro* pull-down assay. Purified CqsS protein was incubated with purified WT CqsA protein, supplemented with Luria-Bertani (LB) medium (lanes 3 and 8) or 10% (v/v) cell-free culture medium from *E. coli* expressing WT CqsA (lanes 5 and 10) or the catalytically-inactive CqsA K240A mutant protein (lanes 4 and 9), followed by affinity purification using the Strep tag on CqsS. The input (lanes 1-5) and output (lanes 6-10) of the pull-down assay were subjected to SDS-PAGE analysis.
