## Extended data Fig. 1-10; Supplementary Table 1; Supplementary Video 1-2 for "Molecular basis of quorum-sensing signal transduction by CqsS and its inhibition by CqsA the autoinducer synthase": Supplementary information.pdf

**Supplementary Table 1. Summary of data collection and model statistics.**

| | CqsS$_{S1}$ EMD-66349 PDB 9WXL | CqsS$_{S2}$ EMD-66354 PDB 9WXQ | CqsS-ATPγS$_{S1}$ EMD-66361 PDB 9WY6 | CqsS-ATPγS$_{S2}$ EMD-66365 PDB 9WY7 | CqsS-CqsA EMD-66360 PDB 9WY4 | CqsS-CAI-1 EMD-66357 PDB 9WXU |
|---|---|---|---|---|---|---|
| **Data collection and processing** | | | | | | |
| EM | Titan Krios (Thermo Fisher) | | | | | |
| Voltage (kV) | 300 | | | | | |
| Detector | BioQutum K3 (Gatan) | | | | | |
| Electron dose ($e^-$/Å$^2$) | 50 | | | | | |
| Pixel size (Å) | 0.82 | | | | | |
| Micrographs | 1,630 | | 1,434 | | 2,731 | |
| **Reconstruction** | | | | | | |
| Software | cryoSPARC | | | | | |
| Particles | 124,184 | 137,177 | 150,052 | 140,065 | 130,780 | 131,035 |
| Symmetry | C1 | | | | | |
| Resolution (Å) | 3.2 | 3.3 | 3.5 | 3.5 | 2.9 | 3.4 |
| Map sharpening B factor (Å$^2$) | 120 | 126 | 136 | 139 | 75 | 162 |
| **Model building, Refinement and Validation** | | | | | | |
| Software | Coot, Phenix | | | | | |
| Model composition | | | | | | |
| Protein residues | 1,149 | 1,030 | 1,057 | 1,045 | 2,500 | 336 |
| Nonhydrogen atoms | 8,711 | 7,932 | 8,207 | 8,046 | 19,091 | 2,904 |
| Ligands | 0 | 0 | 1 | 1 | 4 | 2 |
| R.m.s. deviations | | | | | | |
| Bond lengths (Å) | 0.005 | 0.011 | 0.005 | 0.004 | 0.005 | 0.005 |
| Bond angles (°) | 0.84 | 1.12 | 0.79 | 0.80 | 0.76 | 0.95 |
| MolProbity score | 1.80 | 1.81 | 1.91 | 2.21 | 2.28 | 1.99 |
| Clashscore | 7.69 | 9.96 | 8.59 | 11.44 | 9.60 | 10.67 |
| Poor rotamers (%) | 0.60 | 0.64 | 1.11 | 1.78 | 3.63 | 0.67 |
| Ramachandran plot | | | | | | |
| Favored (%) | 94.35 | 95.87 | 93.89 | 93.13 | 95.01 | 93.03 |
| Allowed (%) | 5.38 | 3.94 | 5.83 | 6.39 | 4.71 | 6.97 |
| Disallowed (%) | 0.26 | 0.20 | 0.29 | 0.48 | 0.28 | 0 |

**Supplementary Video 1. Conformational changes in CqsS from CqsS-ATPγS$_{S1}$, through CqsS$_{S2}$, CqsS-ATPγS$_{S2}$, to CqsS$_{S1}$.**

The video was generated with the CqsS$_{S1}$, CqsS$_{S2}$, CqsS-ATPγS$_{S1}$, and CqsS-ATPγS$_{S2}$ structures.

**Supplementary Video 2. Conformational changes in the TMD between the CqsS apo- and CAI-bound states.**

The video was generated with the structures of CqsS$_{S1}$ and CqsS-CAI-1 as the first and final frames, respectively.
